# Organization and Plasticity of Inhibition in Hippocampal Recurrent Circuits

**DOI:** 10.1101/2023.03.13.532296

**Authors:** Bert Vancura, Tristan Geiller, Attila Losonczy

## Abstract

Excitatory-inhibitory interactions structure recurrent network dynamics for efficient cortical computations. In the CA3 area of the hippocampus, recurrent circuit dynamics, including experience-induced plasticity at excitatory synapses, are thought to play a key role in episodic memory encoding and consolidation via rapid generation and flexible selection of neural ensembles. However, *in vivo* activity of identified inhibitory motifs supporting this recurrent circuitry has remained largely inaccessible, and it is unknown whether CA3 inhibition is also modifiable upon experience. Here we use large-scale, 3-dimensional calcium imaging and retrospective molecular identification in the mouse hippocampus to obtain the first comprehensive description of molecularly-identified CA3 interneuron dynamics during both spatial navigation and sharp-wave ripple (SWR)-associated memory consolidation. Our results uncover subtype-specific dynamics during behaviorally distinct brain-states. Our data also demonstrate predictive, reflective, and experience-driven plastic recruitment of specific inhibitory motifs during SWR-related memory reactivation. Together these results assign active roles for inhibitory circuits in coordinating operations and plasticity in hippocampal recurrent circuits.

## Introduction

Episodic memories formed from single experiences can be used to guide behavior throughout the lifetime of an organism (Tulving, 2002). For this to occur, continuous streams of sensory information must be discretized into snapshots which can then be reactivated and consolidated into long-term memory (Buzsáki, 1989, 2015). In the mammalian brain, both the rapid encoding of episodic memories and their subsequent consolidation rely critically on the CA3 recurrent network in the hippocampus (Daumas et al., 2005; Kesner, 2007; Nakashiba et al., 2009; Nakazawa et al., 2003; Wagatsuma et al., 2017). The initial encoding of episodic memories is thought to be implemented via excitatory synaptic plasticity at synapses between feedforward mossy fibers on CA3 pyramidal cells, as well as at recurrent synapses between pyramidal cells (Mishra et al., 2016; Nakazawa et al., 2002; Rebola et al., 2017). These nascent representations are subsequently consolidated during sharp-wave ripples (SWRs), a synchronous population event generated within the CA3 recurrent network during which a compressed version of the memory traces is thought to be reactivated and transferred to the neocortex for long-term storage (Buzsáki, 2015; Foster, 2017; Gillespie et al., 2021; Norimoto et al., 2018; Pfeiffer, 2020; Wang and Morris, 2010). Accordingly, the main body of work so far has primarily focused on investigating the temporally structured recruitment of CA3 pyramidal cells during memory formation and consolidation (Buzsáki, 2015; Csicsvari et al., 2000; Guzman et al., 2016; Hunt et al., 2018; Leutgeb et al., 2007; Neunuebel and Knierim, 2014), leaving a major question mark regarding the role played by inhibition in supporting and regulating these memory processes (Buzsáki, 2015; Joo and Frank, 2018; Karlsson and Frank, 2009). Indeed, experimental and computational studies strongly implicate inhibitory motifs in stabilizing recurrent networks and supporting efficient cortical computations (Geiller et al., 2021; Nicola and Clopath, 2019; Sadeh and Clopath, 2021; Vogels et al., 2011). However, as the CA1 output region of hippocampus has traditionally served as a prototype circuit for the study of interneurons (Arriaga and Han, 2019; Booker and Vida, 2018; Buzsáki, 2015; Dudok et al., 2021a; Geiller et al., 2020; Klausberger and Somogyi, 2008; Lovett-Barron et al., 2012; McKenzie, 2018; Pelkey et al., 2017; Rebola et al., 2017; Royer et al., 2012) strikingly little is known about inhibitory microcircuits in the upstream recurrent CA3 network and their role in structuring temporally ordered neuronal firing during behaviorally relevant network states associated with memory encoding and consolidation. Based on investigations in CA1, inhibition is predominantly viewed as an immutable pacemaker of principal cell firing, without itself being plastic, a concept reflected in the term ‘chronocircuit’ (Klausberger and Somogyi, 2008). In contrast to this static view of inhibition, *in vivo* structural and molecular studies have revealed robust changes in CA3 inhibitory circuits in response to behavioral manipulations (Donato et al., 2013, 2015; Guo et al., 2018; Ruediger et al., 2011), raising the possibility that CA3 inhibitory dynamics are modifiable in an experience-dependent manner (Rebola et al., 2017; Sharma et al., 2020). Critically, major challenges associated with obtaining population-level recordings of molecularly defined cell types in deep brain regions have impeded addressing these major knowledge gaps concerning operations and plasticity of inhibitory circuits in the CA3 network.

Here, we utilize a two-step method to simultaneously record large numbers of CA3 interneurons with fast, targeted 3-dimensional calcium imaging during behavior and retrospectively identify their molecular profiles with *post hoc* immunohistochemistry. We provide the first comprehensive description of molecularly-identified CA3 interneuron dynamics during both active spatial navigation and awake SWRs associated with memory consolidation. Our results uncover subtype-specific dynamics during behaviorally distinct brain-states, including activity patterns that are both predictive and reflective of SWR properties. Lastly, we provide the first demonstration of subtype-specific, experience-dependent changes in interneuron dynamics around SWRs, suggesting a central role of local inhibition in regulating CA3 microcircuit operations supporting memory consolidation.

## Results

### Large-scale imaging of molecularly identified interneurons

To obtain large-scale recordings from CA3 inhibitory circuits, we injected VGAT-Cre mice with a rAAV encoding Cre-dependent GCaMP7f into CA3 and implanted an imaging window over the alveus of the anterior hippocampus, providing unbiased access to GABAergic interneurons across all CA3 sublayers (Fig 1A). We used acousto-optic deflection (AOD)-based 2p imaging (Geiller et al., 2020; Katona et al., 2012; Szalay et al., 2016) to record from hundreds of interneurons scattered in three dimensions while mice were engaged head-fixed on a treadmill(Geiller et al., 2017) in a spatial foraging task (total of 1,311 interneurons in 14 mice; 93.6 ± 21.6 per mouse, mean ± SD) (Fig 1B, 1C). After the conclusion of imaging experiments, multiplexed *post hoc* immunohistochemistry was performed on fixed brain slices that were registered to *in vivo* structural scans, revealing the molecular identity of imaged cells (Fig 1D, Methods). The precise location of all imaged cells was determined with calbindin (CB) immunohistochemistry, which allows visualization of granule cell mossy fibers (Fig 1E) within the *stratum lucidum* (Dudek et al., 2016). We utilized a combination of five molecular markers (parvalbumin, somatostatin, SATB1, cholecystokinin, and calbindin) to identify five subtypes of CA3 pyramidal cell-targeting interneurons: parvalbumin-positive basket cells (PVBCs), axo-axonic cells (AACs), somatostatin-positive cells (SOMs), cholecystokinin-positive cells (CCKs), and calbindin-positive cells (CBs) (Fig 1F-H, Methods). These five markers were chosen to label some of the most abundant interneuron subtypes in CA3 (Freund and Buzsáki, 1996; Pelkey et al., 2017), including those not readily accessible *via* traditional transgenic driver lines.

**Figure 1.**
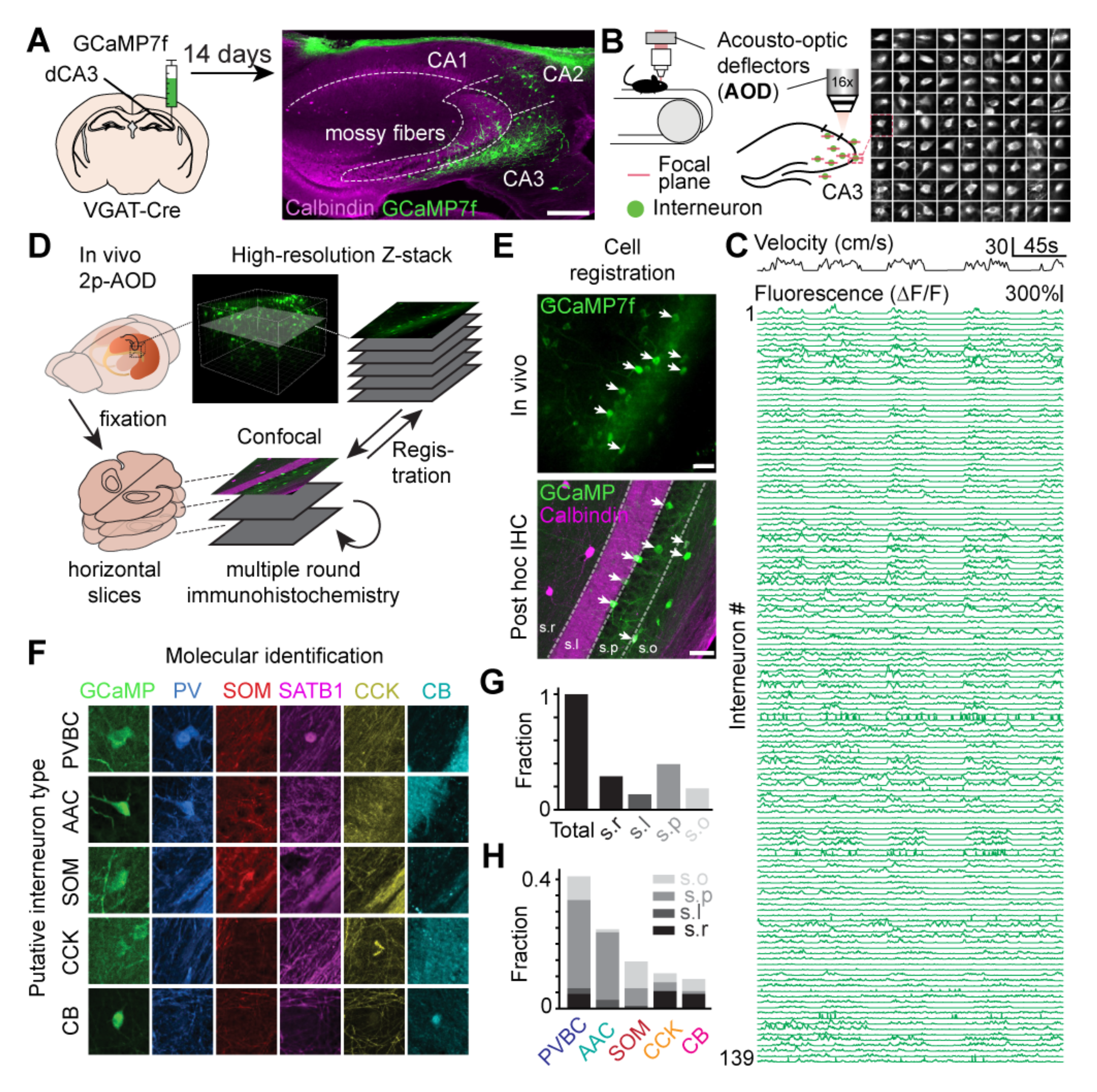
Large-scale imaging of molecularly-identified GABAergic interneurons in CA3. A. Experimental design: VGAT-Cre mice are injected in CA2/CA3 with a Cre-dependent GCaMP7f virus to record all inhibitory interneurons with 2p imaging. Scale bar on the right confocal image represents 250 µm. B. Hundreds of interneurons can be recorded simultaneously at 5-10 Hz in three dimensions with the AOD microscope during behavior. Right: time-average examples of 81 interneurons. The images are 50×50µm. C. Example fluorescence traces from 139 simultaneously recorded interneurons during several minutes of behavior (animal velocity plotted above). D. Schematic of the experimental pipeline used to determine the molecular identity of imaged cells. Multiple rounds of immunohistochemistry were performed on fixed, horizontal slices that were registered to high-resolution *in vivo* Z-stacks. E. Example *in vivo* AOD-2p image (top) and confocal image (bottom) of the registered FOV. White arrows indicate the registered cells. Calbindin immunohistochemistry was used to label the mossy fibers of *stratum lucidum*. Scale bars on the top and bottom images represent 50 and 100 µm, respectively. F. Example immunohistochemical labeling and combinatorial expression patterns of the 5 markers (PV, SOM, SATB1, CCK, CB) used to separate imaged cells into subtypes. All images are approximately 60 × 60 µm. G, H. Layer and subtype distributions of all imaged and *post hoc* identified interneurons.

### Online inhibitory dynamics during spatial navigation

Locomotory movements are the behavioral correlates of an online and actively engaged brain-state, characterized in the hippocampus by a location-specific rate code in pyramidal cells. Thus, locomotion has been shown to strongly influence the recruitment or disengagement of distinct types of interneurons in CA1 (Arriaga and Han, 2017; Dudok et al., 2021b), but the paucity of data in CA3 to date has hindered efforts in testing whether these dynamics are a global interneuron signature hippocampus-wide. We trained and imaged water-restricted mice during a random foraging task on a 2m-long cue-rich belt (Geiller et al., 2020), during which several water rewards were delivered at random locations on each lap (Fig 2A). We observed a tight correlation between the activity of most cells and the ambulatory state of the animal (locomotion vs. immobility) (Fig 2B). To systematically characterize this relationship, we calculated Pearson’s correlation coefficient between the activity of each cell and the animal’s velocity. At the population level, the average correlation was shifted towards positive values (Fig 2C). We then compared the coefficients across subtypes and observed AACs to be overall more tightly correlated with velocity (Fig 2D). In addition, CB cells had overall lower coefficients, and many CB cells were more active during immobility than during locomotion (Fig 2D). Closer inspection of the immobility-active CB cells revealed the vast majority lacked SATB1 expression (CB+/SATB1-), while SATB1-positive CB cells (CB+/SATB1+) displayed high correlations with locomotion (Fig 2E-G). These subtype-specific trends were also reflected in the average run-start and run-stop responses for each subtype (Supp Figure 1). As these functionally unique CB+/SATB1- cells may represent a previously unrecognized interneuron subtype and were negative for the other tested markers (PV, SOM, CCK), we next determined what other molecules they may express and found that the majority of CB cells were also negative for most other hippocampal interneuron markers (VIP, CR, NPY, M2R) and expressed the transcription factor COUP-TFII (Supp Figure 2), although no differences were seen between SATB1+ and SATB1- cells.

**Figure 2.**
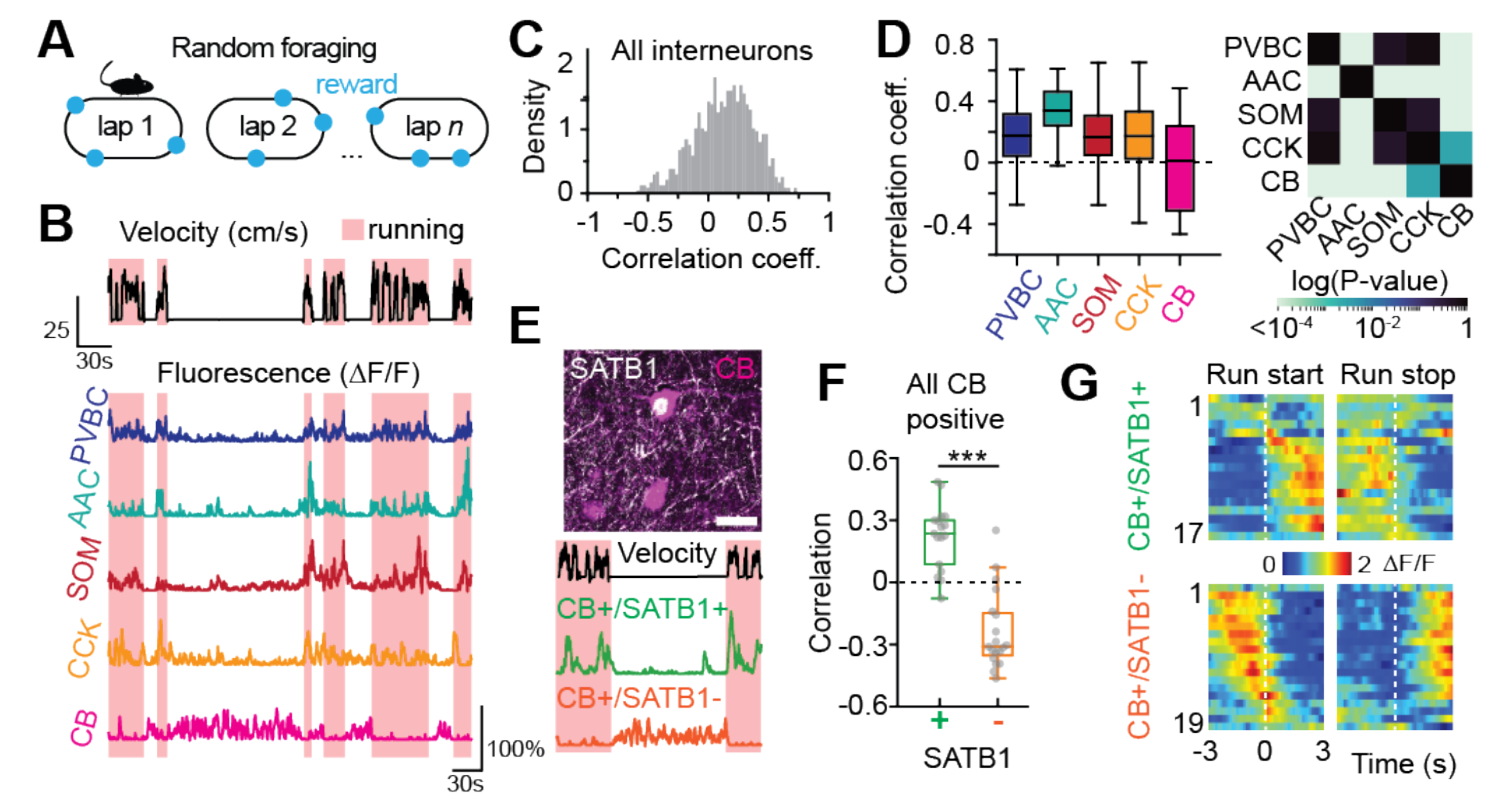
Online dynamics during spatial navigation. A. Mice are trained to run for randomly delivered water rewards on each lap. B. Representative fluorescence traces from different interneuron subtypes during running (red shaded area) and immobility (non-shaded) bouts. C. Distribution of Pearson’s correlation coefficients between fluorescence and velocity for all recorded interneurons. D. Same as C for all identified subtypes. Between-subtype statistical comparisons (t-test) are represented in the heatmap on the right (n = 121 PVBCs, 89 AACs, 86 SOMs, 67 CCKs, and 36 CBs from n = 9 mice). E. *Top:* Confocal image of a CB+/SATB1+ cell (magenta/white) and a CB+/SATB1- cell (magenta only). Scale bar represents 25 µm. *Bottom:* Example fluorescence traces from these cells during locomotion and immobility. F. Same as D for CB+/SATB1+ and CB+/SATB1- subtypes (n = 17 CB+/SATB1+ neurons; n = 19 CB+/SATB1- neurons, from n = 9 mice). CB+/SATB1+ cells were significantly more correlated with velocity than CB+/SATB1- cells (Mann-Whitney U Test, p = 2.17 × 10^-6^). G. Heatmaps of average activity around run-start (left) and run-stop (right) events for all CB+/SATB1+ (top) and CB+/SATB1- (bottom) interneurons, sorted by the location of their peak activity around run-start events (the same row on the left and right heatmaps represents the same cell).

Akin to pyramidal cells selective for particular regions of an environment (O’Keefe and Dostrovsky, 1971), interneurons have also been reported to display some degree of spatial selectivity in CA1 (Ego-Stengel and Wilson, 2007; Geiller et al., 2020; Grienberger et al., 2017; Hangya et al., 2010; Marshall et al., 2002; Wilent and Nitz, 2007). Thus, we next sought to determine the nature and extent of spatial tuning by quantifying the degree of selectivity among different CA3 interneuron subtypes (Supp Figure 3). While subsets of cells displayed both significant positive and negative spatial tuning, we did not observe profound differences between subtypes (Supp Figure 3A-D). We then more precisely quantified the relative contributions of various behavioral variables to interneuron activity by constructing a model to predict each cell’s fluorescence trace (Supp Figure 3E-I) and found that velocity, followed by position, displayed the highest predictive coefficient.

### Inhibitory dynamics around offline memory consolidation events

The precise circuit mechanisms regulating SWR initiation in CA3 and the role played by distinct inhibitory interneurons in these processes remain unknown, as no data exists from defined interneuron subtypes. To address this major knowledge gap, we used our 2p- AOD method combined with local field potential (LFP) recordings (Fig 3A, Methods) from contralateral CA1 to examine interneuron activity around awake SWRs during periods of immobility. To confirm that we could detect changes in CA3 circuits around contralateral SWR events (Grosmark et al., 2021; Guan et al., 2021; Kohl et al., 2011; Malvache et al., 2016; Terada et al., 2021),we first imaged CA3 pyramidal cell activity using a CA3-specific transgenic mouse line (Supp Figure 5A, B). CA3 pyramidal cell activity was indeed elevated around detected SWRs (Supp Figure 5C), allowing us to next consider the local inhibitory dynamics. At the overall population level and regardless of subtype, we found that the first and second components of a principal component analysis-based decomposition represented activated and inhibited dynamics, respectively, after and around SWR onset (Fig 3B). In stark contrast with the available *in vitro* data where most CA3 interneurons spike during SWRs (Hájos et al., 2013), we in fact found more cells to be inhibited (Fig 3B). We observed highly heterogeneous responses at the subtype level as evidenced by their average peri-SWR traces (Fig 3C, D): PVBCs showed a tight time-locked activation around SWR events, while AACs, CCKs and CB cells showed a net inhibition. CCKs were unique in that the trough of their inhibition preceded the SWR onset (Fig 3C). We then examined how representative these average responses were for all cells belonging to each subtype. To this end, we calculated a SWR activity index to measure whether a given interneuron is more activated or inhibited during SWRs (Fig 3E). While most PVBCs were strongly activated, the majority of AACs, CCKs, and CBs were inhibited (Fig 3E, F). SOM cells exhibited a bimodal profile, including the most activated cells in the entire dataset but also strongly suppressed ones, explaining the weakly modulated overall peri-SWR trace of the SOM subtype (Fig 3C, E, F and Supp Figure 4); this heterogeneity likely reflects the presence of distinct SOM-expressing subtypes not captured by our molecular identification (Katona et al., 2014; Pelkey et al., 2017). Lastly, we analyzed the timing of individual cells’ responses by examining the location of the peaks of activated cells and the troughs of inhibited cells around SWRs, depending on whether a given subtype is composed of a larger fraction of activated or inhibited interneurons (‘dominant response’) (Fig 3G). While most inhibited cells tended to have their troughs well after the SWR, inhibited CCK cells had their trough before the initiation of SWRs (Fig 3G), ideally positioning them to exert control over the initiation of SWRs bound to the synchronous activity of CA3 pyramidal cells (Fig 3G).

**Figure 3.**
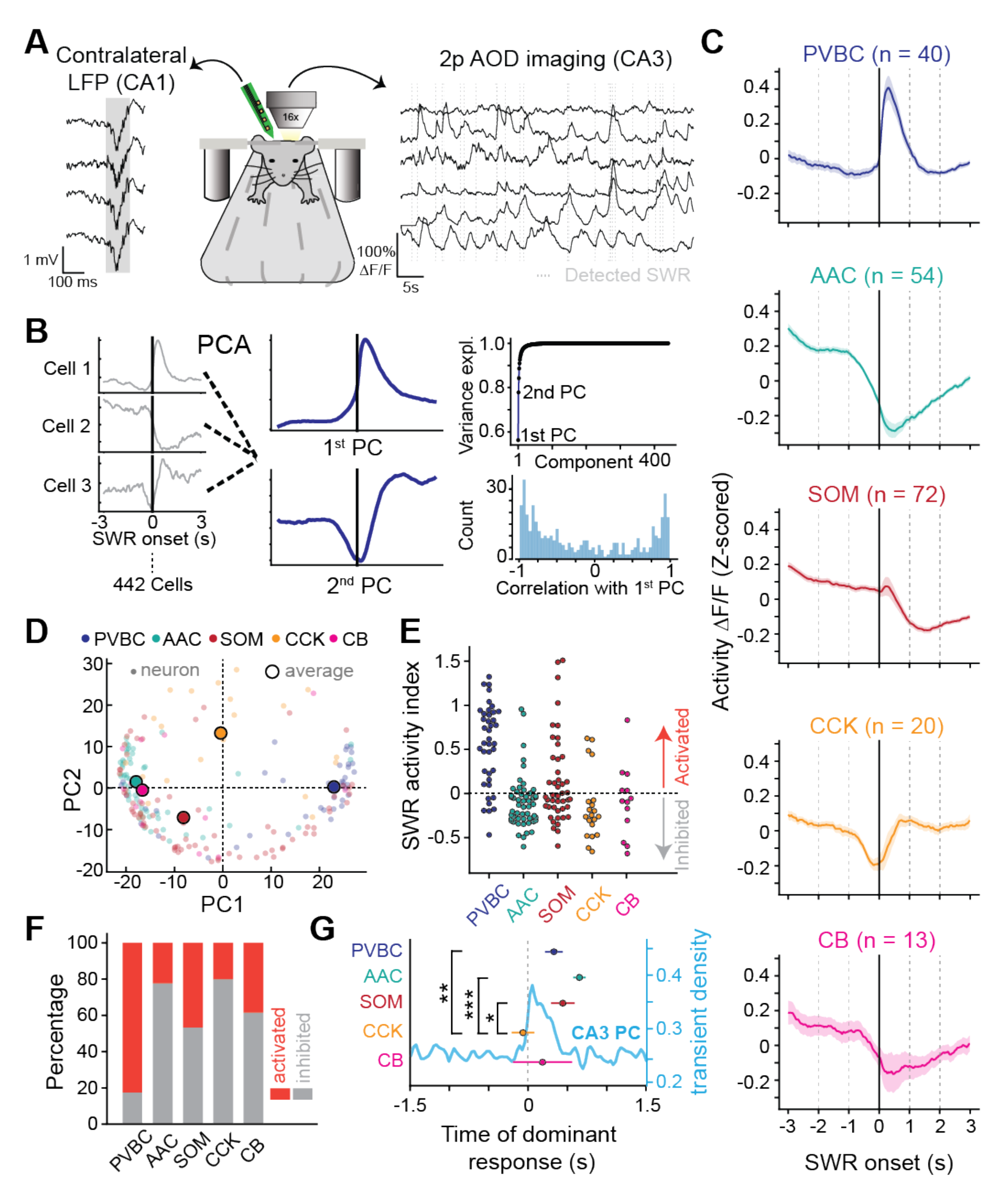
Subtype-specific offline dynamics during SWR events. A. Experimental setup for simultaneous 2p-AOD imaging and LFP recordings. SWRs were recorded on a 4-channel silicon probe implanted in the contralateral CA1. B. PCA was performed on the average peri-SWR traces of all cells (left) to produce the first and second principal component time series (center) (n = 442 cells from n = 5 mice). Cumulative explained variance with each additional principal component (top right). Histogram of the correlation coefficient of each cell’s average peri-SWR trace with the first principal component (bottom right). C. Average peri-SWR traces for all subtypes (data from n = 5 mice). D. Distribution of first and second PC loadings for all cells (small gray dots) and the median values for all identified subtypes (larger colored dots, n = 40 PVBCs, 54 AACs, 72 SOMs, 20 CCKs, and 13 CBs from n = 5 mice). E. Average SWR activity index for all neurons, grouped by subtype (number of identified cells of each subtype and number of mice is the same as above). As a whole, PVBCs were significantly activated, AACs were significantly inhibited, SOMs were weakly activated, and CCKs and CBs showed no overall net modulation (one-sample t-tests against 0: PVBC: p = 5.52 × 10^-8^; AAC: p = 2.39 × 10^-6^; SOM: p = 0.028; CCK: p = 0.99; CB: p = 0.57). F. Fraction of inhibited and activated neurons for each subtype, defined by the SWR activity index. G. Timing of the dominant response for each subtype as a function of SWR onset. Dominant response was characterized by the highest fraction of neurons within a given subtype (PVBC’s dominant response is activated while AAC is inhibited). The corresponding peak or trough location was then calculated for all the neurons falling in the dominant’s category (unpaired t-test, p(CCK-SOM)=0.02, p(CCK- AAC)=2.2 × 10^-5^, p(CCK-PVBC)=5.8 × 10^-3^).

### Predictive and reflective inhibitory dynamics around SWRs

SWR power recorded *in vivo* exhibits marked event-to-event variability which is thought to reflect differences in the size of recruited pyramidal cell ensembles in the hippocampal network (Csicsvari et al., 2000; Buzsáki, 2015; Grosmark and Buzsáki, 2016). If inhibitory subtypes influence SWR initiation or termination in the CA3 network, we expect that the activity dynamics of those cells would be correlated with the SWR power. To test this, we first sought to confirm that we could detect differences in local CA3 pyramidal cell network dynamics during high- and low-power SWRs measured on the contralateral side with our imaging approach. As the pyramidal cell transient rate was significantly greater around high-power SWRs than around low-power SWRs (Supp Figure 5D, E), we next sought to determine whether similar correlations could be observed between interneuron dynamics and SWR power. We found that some interneurons indeed displayed responses largely correlated with the amplitude of the SWR they follow (PVBC) or precede (CCK) (Fig 4A). After splitting the dataset between the lowest (0-20^th^ percentile) and the highest (80-100^th^ percentile) SWR power, only PVBC, AAC and CCK subtypes showed a different average response trace (Fig 4B, C). The position in time of the most extreme difference between the average low- and high-SWR responses was, for CCKs, tens of milliseconds before the SWR onset, while AACs and PVBCs had differential responses after SWR onset (Fig 4D). These results suggest that CCK activity before a given SWR should be significantly more predictive of the resulting SWR power than the activity of the other subtypes, while PVBC and AAC activity after a given SWR should be selectively more reflective of the SWR power. To test this more directly, we correlated the average activity of each cell in the 500 ms before and after each SWR with the magnitude of the SWR (Methods, Fig 4E, F). Indeed, we found significant subtype-specific negative and positive relationships: large amplitude SWRs were associated with decreased CCK activity preceding the event, as well as increased PVBC activity and decreased AAC activity following it (Fig 4G). Overall, these results suggest a novel role for CCK interneurons in controlling the size of the population burst that generates the SWR, as well as an important role for PVBCs in regulating peri-SWR network activity.

**Figure 4.**
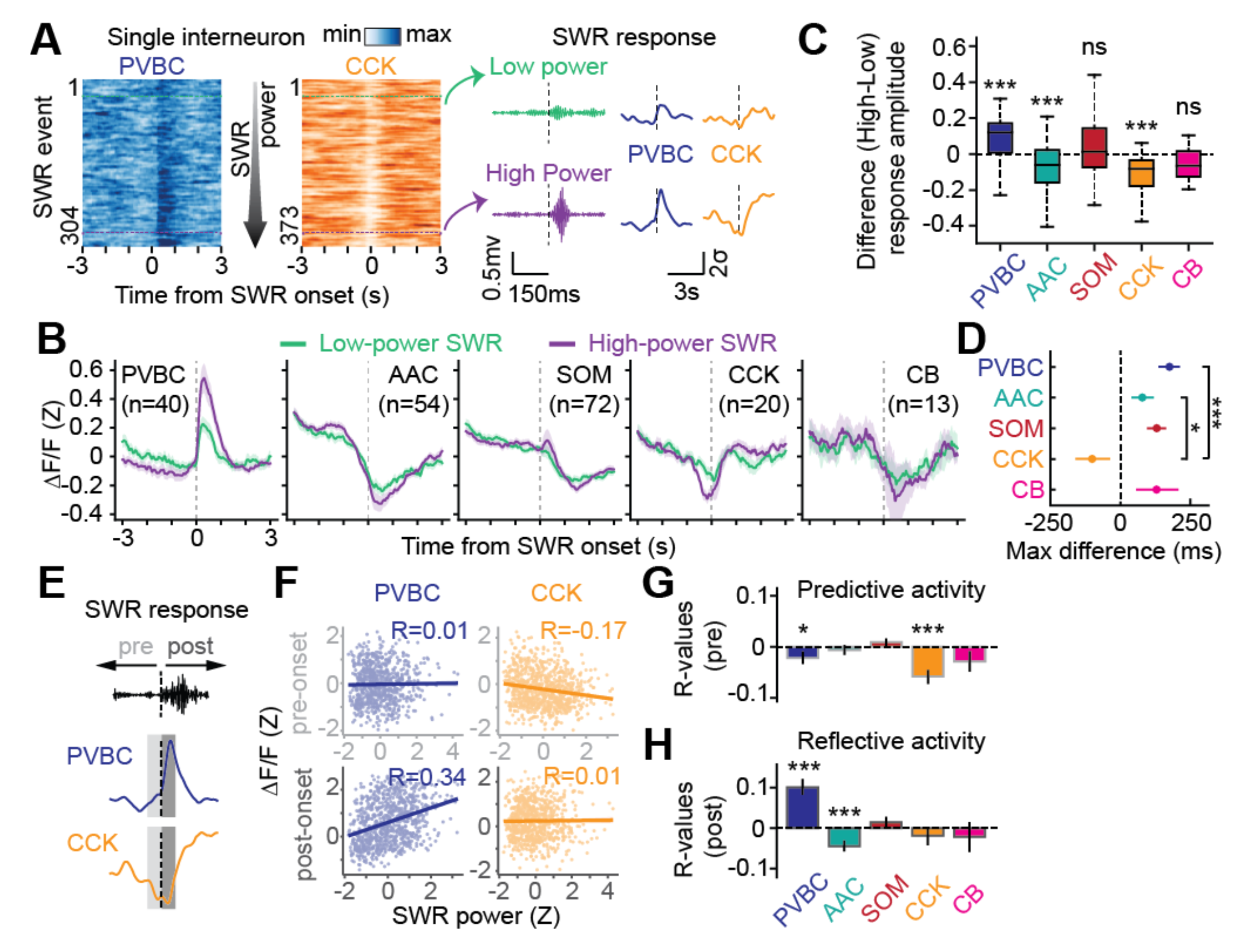
Peri-SWR dynamics are both predictive and reflective of SWR properties in a subtype-specific manner. A. Example responses of a PVBC and a CCK cell to high- and low-power SWRs. Heatmaps illustrating cell responses around SWR events ordered by their power (left). Both activated and inhibited responses were strongly modulated by SWR power (right). B. Average Z-scored peri-SWR traces for both low- (0-20^th^ percentile) and high- (80- 100^th^ percentile) power SWRs for each subtype (number of cells of each subtype indicated on the figure, n = 5 mice). C. Average value of the difference in activity between high- and low-power SWRs for each cell, grouped by subtype. PVBCs were significantly more activated around high-power SWRs, while both AACs and CCKs were significantly more inhibited around high-power SWRs (n = 39 PVBCs, 53 AACs, 72 SOMs, 20 CCKs, and 13 CBs from n = 5 mice; one-sample t-tests against 0: PVBC: p = 2.05 × 10^-4^; AAC: p = 8.16 × 10^-5^; SOM: p = 0.11; CCK: p = 5.76 × 10^-4^; CB: p = 0.24). D. Location of the maximum difference in activity between high- and low-power SWRs for each cell, grouped by subtype. CCKs were most prominently modulated by high-amplitude SWRs earlier in time compared to AACs and PVBCs, the other two modulated subtypes (same number of cells and mice as in C; Mann-Whitney U Tests, p (CCK-AAC) = 0.015; p (CCK-PVBC) = 5.38 × 10^-4^). SOM and CB cells, while not on average differentially modulated by high- vs. low-amplitude SWRs, are shown here for visual comparison. E. Schematic illustrating the pre- and post-activity around each SWR used for the correlation analysis. A 500 ms window before and after each SWR event was used. F. Example correlation plots between either average pre- (top) or average post-activity (bottom) and SWR power for individual PVBC (left) and CCK cells (right). Correlations include every SWR event during which the cell was imaged and generate an r-value for each cell. G. Summary of r-values for all cells for the correlation between each cell’s average pre-SWR activity and SWR power, grouped by subtype. CCK activity was most predictive of SWR power (lower CCK activity before high-power SWRs), followed by PVBC activity. The other subtypes were not predictive of SWR power (same number of cells as in C, data from n = 5 mice; one-sample t-tests against 0: PVBC: p = 0.033; AAC: p = 0.47; SOM: p = 0.10; CCK: p = 1.72 × 10^-4^; CB: p = 0.16). H. Same plot as in G, but now for the correlation between each cell’s average post-SWR activity and SWR power, grouped by subtype. PVBC and AAC activity were both reflective of SWR power (higher PVBC and lower AAC activity following high-power SWRs). The other subtypes were not reflective of SWR power (same number of cells as in C, data from n = 5 mice; one-sample t-tests against 0: PVBC: p = 1.16 × 10^-6^; AAC: p = 1.14 × 10^-4^; SOM: p = 0.22; CCK: p = 0.39; CB: p = 0.56).

### Experience-dependent reconfiguration of peri-SWR inhibitory dynamics

Selective recruitment and exclusion of CA3 pyramidal cells during SWRs depending on behavioral relevance of stimuli they encode represent important mechanisms for selective consolidation of salient representations (Terada et al., 2017). While these phenomena suggest a role for inhibitory plasticity, circuit mechanisms enabling the flexible recruitment of individual pyramidal cells to SWRs remain unknown. Instead, interneurons are thought to have stereotyped and immutable activity profiles around SWR events that are subtype-specific (Klausberger and Somogyi, 2008; Somogyi et al., 2014; Varga et al., 2012). Therefore, potential behavior-dependent changes in recruitment of interneurons to SWR events, and the resulting flexible selection of pyramidal cell assemblies to these events, remains an open question. To directly address this, we leveraged a recent finding that CA3 pyramidal cells encoding sensory cues are suppressed from SWRs when the cues have no behavioral relevance (Terada et al., 2017). Thus, we asked whether changes in the activity of CA3 interneurons around SWRs occur immediately after sensory stimulation (Fig 5D). To this end, mice received pseudorandomly delivered water, light, and odor stimuli while head-fixed on a cue-less burlap belt(Terada et al., 2021), and interneurons were imaged during SWRs before and after the sensory stimulation (Fig 5A). CCK cells were selectively activated in response to the sensory stimuli (Fig 5B). An activation could be seen in the vast majority of CCK cells, the response did not change significantly over trials for any subtype, and the response was generally similar for the three sensory modalities (Fig 5C, Supp Fig 6A, B, C). Next, we constructed average peri-SWR traces for each subtype, both before and after sensory stimulation (Fig 5E, Supp Figure 6D). While most subtypes (PVBC, AAC, and SOM cells) displayed stable peri-SWR activity profiles, CCK cells showed a significantly increased activation around SWR events (Fig 5E) after the sensory stimulation. Additionally, several CB cells displayed a similar increase in peri-SWR activity, although the change was not significant at the subtype level. We quantified this change at the level of individual cells, calculating the peri-SWR modulation for each cell both before and after sensory stimulation (Fig 5F, Methods). This analysis revealed a selective increase in SWR modulation for CCK cells in response to sensory stimulation (Fig 5F). We next asked whether this increase in the overall peri-SWR activity of CCK cells was due to an increased fraction of events recruited to, or to an increased activation in a comparative number of SWR events. We found that individual CCK cells were recruited to an increased number of SWR events post sensory stimulation, while the maximum activation within recruited events remained stable (Fig 5G, 5H). Together, these results provide the first evidence for subtype-specific changes in interneuron recruitment to SWR events in response to experience and suggest a potential role for CCK interneurons in the regulation of the ensemble size and identity of pyramidal cells recruited to SWRs.

**Fig 5.**
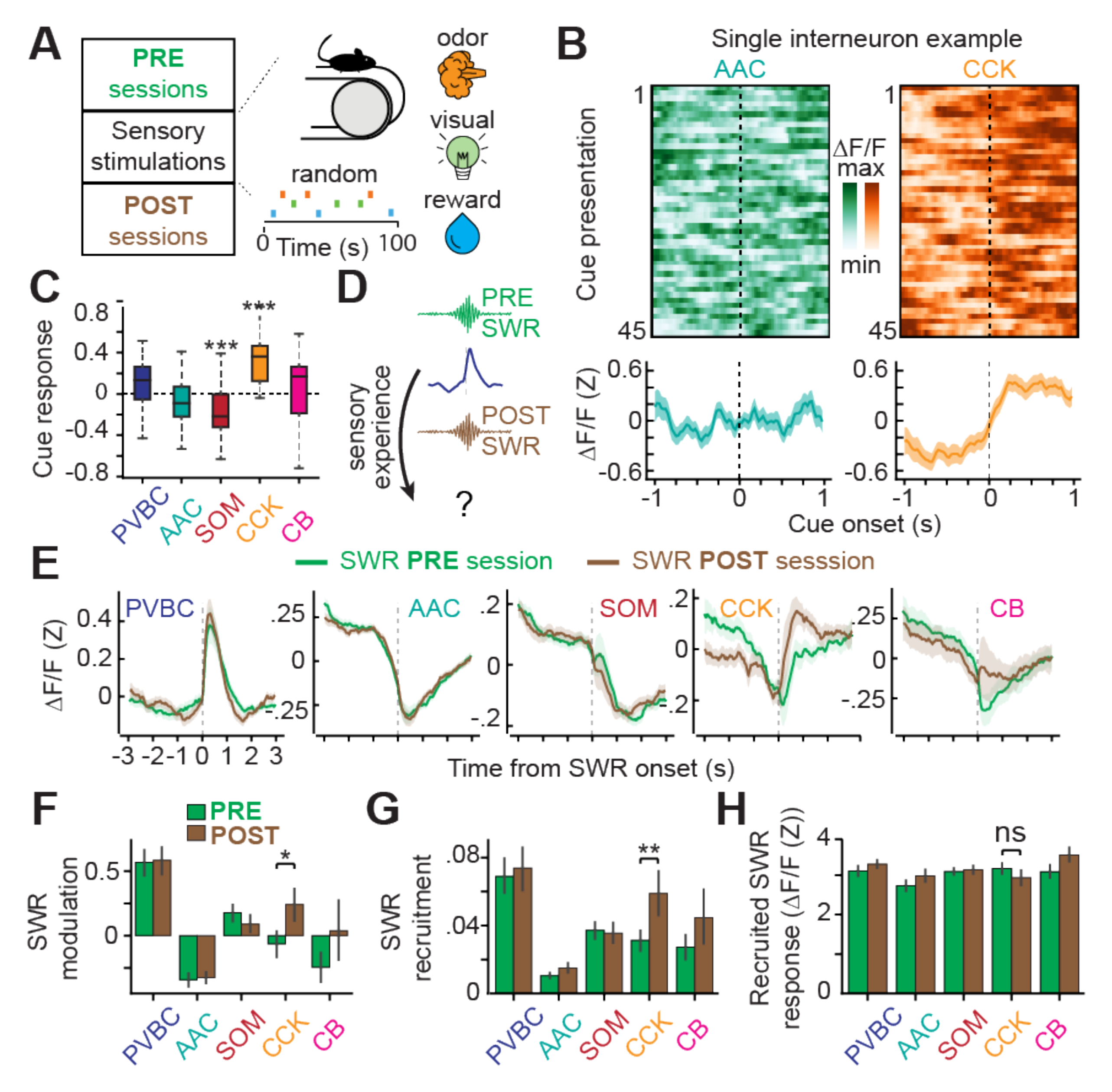
Peri-SWR dynamics can be modulated by experience. A. Sensory stimulation paradigm. Water, light, and odor stimuli were presented pseudorandomly while the mouse remained head-fixed on a cue-less, burlap belt. Interneurons were imaged during SWRs in both the PRE and POST sessions as well as during stimulus presentations. B. Representative example of an individual AAC (green) and CCK (orange) interneuron. Heatmaps represent the activity during all sensory stimulus presentations (45 in total) with the corresponding average response (bottom). The CCK neuron is consistently and significantly activated by cue presentations. C. Average sensory cue response for each cell, grouped by subtype. CCK cells were significantly activated by cue presentation, while SOM cells were significantly inhibited (n = 25 PVBCs, 37 AACs, 42 SOMs, 14 CCKs, and 12 CBs from n = 3 mice; one-sample t-tests against 0: PVBC: p = 0.076; AAC: p = 0.059; SOM: p = 1.01 × 10^-4^; CCK: p = 1.65 × 10^-4^; CB: p = 0.73). D. Sessions PRE and POST cue presentations are split to examine whether sensory cue presentations induced a change in interneuron dynamics around SWRs. E. Average Z-scored peri-SWR traces for both PRE and POST sessions for all subtypes. Note the different peri-SWR dynamics for CCK cells and the relative stability of the other subtypes. F. Average SWR modulation for each cell during both the PRE and POST sessions, grouped by subtype. CCK cells had a significantly greater SWR modulation during the POST sessions compared to PRE (n = 29 PVBCs, 35 AACs, 60 SOMs, 16 CCKs, and 9 CBs from n = 5 mice; Wilcoxon signed-rank tests: PVBC: p = 0.97; AAC: p = 0.50, SOM: p = 0.10, CCK: p = 0.013; CB: p = 0.21). G. Fraction of SWR events each cell was recruited to for all cells during both the PRE and POST sessions, grouped by subtype. CCK cells were recruited to a significantly greater fraction of SWR events during the POST session compared to PRE (same number of cells and mice as in F; Wilcoxon signed-rank tests: PVBC: p = 0.75; AAC: p = 0.13; SOM: p = 0.55; CCK: p = 0.0061; CB: p = 0.59). H. Average response magnitude of each cell within recruited SWR events, grouped by subtype. The response magnitude of CCK cells (or any other subtype) within recruited SWRs did not change between PRE and POST (n = 28 PVBC, 29 AACs, 51 SOMs, 13 CCKs, and 8 CBs from n = 5 mice; Wilcoxon signed-rank tests: PVBC: p = 0.37; AAC: p = 0.48; SOM: p = 0.85; CCK: p = 0.17; CB: p = 0.21).

## Discussion

The work here represents the first large-scale assessment of CA3 interneuron dynamics during spatial navigation and SWR-associated memory reactivation, providing critical new information on the activity and plasticity of inhibitory circuits at the locus of SWR initiation. The available data regarding CA3 interneurons largely comes from studies detailing their electrophysiological properties *in vitro* (Botcher et al., 2014; Gulyás et al., 1992; Kohus et al., 2016; Losonczy et al., 2004; Mercer et al., 2007; Papp et al., 2013; Spruston et al., 1997; Vida and Frotscher, 2000), especially as they relate to innervation and plasticity at the mossy fiber – interneuron synapse (Acsády et al., 1998; Lawrence and McBain, 2003; Maccaferri et al., 1998; Mori et al., 2007; Rebola et al., 2017; Szabadics and Soltesz, 2009), and the role of CA3 interneurons in generating SWRs (Bazelot et al., 2016; Dugladze et al., 2012; Ellender et al., 2010; Schlingloff et al., 2014). A complementary line of work has leveraged demanding juxtacellular recordings to record from small numbers of anatomically identified CA3 interneurons under anesthesia *in vivo* (Lasztóczi et al., 2011; Tukker et al., 2013; Viney et al., 2013). Thus, to date, the sparsity, diversity, and depth of CA3 inhibitory circuits in the intact brain have prevented cell-type specific population recordings during behavior from this critical hippocampal region. Our approach combining large-scale, unbiased sampling of inhibitory network activity during behavior, simultaneous LFP recordings, and *post hoc*, multiplexed molecular characterization allowed us to make novel observations regarding the subtype-specific and brain-state-dependent modulation of inhibition in CA3. We note that many of our findings would not have been achieved with more traditional *in vivo* imaging approaches using transgenic lines, highlighting the power of this experimental strategy.

We find that the *in vivo* dynamics of CA3 inhibitory circuits exhibit similarities to their CA1 counterparts with respect to their overall positive velocity modulation and spatial modulated firing in a subset of interneurons. One notable exception is the CCKs that are almost exclusively immobility active in CA1 (Dudok et al., 2021b; Geiller et al., 2020), but more heterogenous in CA3. Our finding that CA3 CB interneurons comprise two functionally distinct subpopulations with regards to their activity during locomotion, and that these subpopulations strongly correlate with expression of the transcription factor SATB1, represents the first *in vivo* characterization of CB interneuron dynamics in any hippocampal subregion. Although these cells represent a significant inhibitory subpopulation (Bezaire and Soltesz, 2013; Freund and Buzsáki, 1996)they have been understudied, likely because of their preferred location in deep dendritic layers (*radiatum/lacunosum-moleculare*), and because major subpopulations of pyramidal cells in CA1 also express CB, complicating genetic access with transgenic lines (Dong et al., 2009). The immobility-active CB cells we identify are potentially linked to a rare subtype previously identified in single-cell transcriptional data from CA1 (Harris et al., 2018) and recorded from in CA1 under anesthesia (Fuentealba et al., 2010). As SATB1 acts downstream of other transcription factors that specify an origin from the medial ganglionic eminence (MGE) and is thus enriched in MGE-derived interneurons (Close et al., 2012; Denaxa et al., 2012), our finding suggests that the functional dichotomy we observe in CB-expressing interneurons may represent differences between MGE- and CGE-derived CB interneurons. We hypothesize that differences in the hard-wired input connectivities onto these two cell types might account for their complementary activity profiles during locomotion and immobility, and that these connectivity differences likely apply to MGE- and CGE-derived interneurons more generally. Indeed, to date all hippocampal interneuron subtypes with robust and consistent immobility activation are CGE-derived, including CCK basket cells (Dudok et al., 2021b), VIP/M2R interneurons (Francavilla et al., 2018), and here CB+/SATB1- interneurons. In support of this view, CCK basket cells and CB-expressing cells in CA1 have both been shown to receive large fractions of inhibitory synapses compared to other interneuron subtypes (Gulyás et al., 1999; Matyas et al., 2004), potentially explaining how these cells could be silenced during locomotion, when the majority of interneurons are activated and would provide them with a strong inhibitory input. Complementarily, a common excitatory input to these cell types that is activated during immobility could explain their sustained immobility activation. Although such an input remains to be found, neuromodulatory afferents are attractive candidates (Atherton et al., 2015; Kaufman et al., 2020; Prince et al., 2021), as they have been shown to regulate interneuron dynamics during locomotion/immobility in the neocortex (Fu et al., 2014). Of particular interest, serotonergic afferents from the median raphe have been shown to selectively target CCK and CB cells in CA1 (Freund et al., 1990; Varga et al., 2009), and certain serotonin receptor subtypes are depolarizing and restricted to CGE-derived interneurons (5-HT3AR). It will be critical in future experiments to understand whether CA3’s CB+/SATB1- and CA1’s VIP+/CCK+ or VIP+/M2R+ neurons share common inputs, if locomotion/immobility signals are passed from one region to the other, and why this locomotion-state dependent inhibition from specific interneuron subtypes is important in regulating information processing in pyramidal cell networks.

While the robust engagement of most hippocampal interneurons is critical for the control of pyramidal cell dynamics during locomotion, spatial navigation, and online memory encoding, their modulation during the SWRs of subsequent immobility is equally crucial for memory consolidation. Our work provides the first data regarding *in vivo* dynamics of defined interneuron subtypes in CA3 around awake SWRs and thus offers important clues regarding the mechanisms of SWR initiation and termination. Although it is now established that SWRs are brain-wide events indispensable for memory reactivation and consolidation, the mechanisms by which they are generated in the hippocampal recurrent network are still under intense debate. While it was originally proposed that SWRs are generated by the disinhibition of recurrently connected CA3 pyramidal cells (Buzsáki, 1986) and more recent CA3 circuit models have also explored this idea (Evangelista et al., 2020), experimental support for this hypothesis has been limited to the finding that a few AACs in CA3 have been found to be silenced during SWRs under anesthesia (Viney et al., 2013). On the contrary, most experiments with *in vitro* CA3 preparations have observed that the vast majority of local interneurons spike during SWRs (Hájos et al., 2013). We find that the majority of AACs and CCKs are inhibited, starting hundreds of milliseconds before the SWR event (see Figure 3), potentially providing the disinhibition necessary to trigger the synchronous activation of pyramidal cells. However, while the AAC inhibition before the SWR is not correlated with the magnitude of the oscillation, the extent of CCK inhibition preceding the SWR is strongly and specifically predictive of the resulting SWR power (see Figure 4). Thus, while AAC inhibition and the resulting withdrawal of inhibition from the axon initial segment of pyramidal cells may be a necessary condition for SWRs to occur, our data suggest that AACs are not responsible for regulating the size of the CA3PC burst. Our results instead ascribe this critical function to CCK-expressing interneurons.

Alternative models of SWR initiation in the CA3 microcircuit posit that the strong activation of local PVBCs may strongly hyperpolarize surrounding pyramidal cells, and the rebound activity from this suppression could generate the CA3 pyramidal cell bursts and the resulting SWR (Buzsáki, 2015). Our results do not support this hypothesis, as the peak of PVBC activation occurs several hundred milliseconds after the SWR event (See Figure 3), and after the peak pyramidal cell response. Nevertheless, we do find that the magnitude of the PVBC response around SWRs is correlated with the magnitude of the oscillation (See Figure 4). This finding extends the *in vitro* CA1 observations that the number of spikes emitted by PV+ cells during sharp waves (SWs) is linearly correlated to the SW amplitude, and that PV+ cells receive EPSCs during SWRs that strongly correlate with SW size (Mizunuma et al., 2014). Relatedly, recent experiments with *in vivo* intracellular recordings from CA3 pyramidal cells during awake SWRs have found that larger SWR events are associated with a more pronounced hyperpolarization in most neurons after the SWR; our finding provides a potential cellular source for this increased inhibition (Kajikawa et al., 2021). In the context of this previous work, our findings suggest that CA3 PVBCs provide feedback inhibition to the activated pyramidal network during SWRs, to structure temporal activity of pyramidal cell ensembles and to preclude the recurrent circuitry from sliding into a degenerated state. Together, our results indicate a complementary role of CCKs and PVBCs (Freund, 2003; Klausberger et al., 2005) in organizing pyramidal cell assembly recruitment in a predictive and reflective manner, respectively.

Our finding of a prominent pre-SWR inhibition in both CA3 AACs and CCKs also raises the critical question: What input silences these cell types, and thus may be responsible for SWR initiation? One attractive candidate is GABAergic afferents from the medial septum. Indeed, subsets of medial septal GABAergic cells have been shown to selectively target AACs and CCKs in CA3 (Joshi et al., 2017), and some medial septal GABAergic cells have been shown to be activated during hippocampal SWRs (Viney et al., 2013). However, many subcortical areas have been shown to be modulated before SWRs (Logothetis et al., 2012), some of which have known afferents to the hippocampus (Atherton et al., 2015; Prince et al., 2021). This also includes subsets of raphe cells (Varga et al., 2009) that decrease their firing rate approximately one second before SWRs (Wang et al., 2015). Furthermore, cholinergic inputs from the medial septum have been shown to regulate SWR rates, as optogenetic activation of these inputs suppresses SWRs (Vandecasteele et al., 2014). Future experiments will be necessary to determine how subcortical neuromodulators interact with defined interneuron subtypes and pyramidal cells in the CA3 network to initiate, propagate, and terminate SWRs, as well as to determine the necessity and sufficiency of each of these microcircuit components in these processes.

Our finding that CCK-expressing interneurons significantly alter their peri-SWR dynamics in response to simple sensory stimulation provides, to the best of our knowledge, the first demonstration of inhibitory plasticity around awake SWRs. It is now well appreciated that pyramidal cell recruitment to SWRs can be modulated based on the animal’s experience, as pyramidal cells encoding behaviorally-relevant stimuli are robustly replayed during SWRs in time-compressed sequences (Grosmark and Buzsáki, 2016; Grosmark et al., 2021; Terada et al., 2021). This dynamic regulation of pyramidal cell recruitment to SWRs is thought to subserve memory consolidation, as behaviorally relevant representations can be preferentially consolidated into long-term memory at the expense of less important ones. In this conceptual framework, inhibition is thought to control the precise timing of pyramidal cell firing but is not itself plastic; our findings challenge this view. We propose that the flexible modulation of CCK-expressing interneuron dynamics around SWRs, and perhaps those of other subtypes not identified in this work, can serve to regulate the flexible recruitment of pyramidal cell subsets to SWRs, profoundly influencing which hippocampal representations are preferentially consolidated into long-term memory. These results are also in agreement with a line of work suggesting that CCKs represent a highly plastic and modifiable motif of hippocampal and neocortical circuits (Hartzell et al., 2018; Klausberger et al., 2005; Del Pino et al., 2017; Yap et al., 2021). Our finding of inhibitory plasticity is especially relevant in CA3, where the recurrent circuitry is ideally suited for the rapid generation and flexible selection of pyramidal cell sequences to SWRs (Nakazawa et al., 2003; Kesner, 2007; Guzman et al., 2016; Rebola, Carta and Mulle, 2017; Nicola and Clopath, 2019). In contrast to the highly plastic recruitment of CCKs, SWR-related activity of other inhibitory motifs (PVBCs, AACs, SOMs) remains unaltered by sensory experience. The stable recruitment of these subtypes can serve to maintain precise temporal and spatial organization (Klausberger and Somogyi, 2008)and stabilization of recurrent network dynamics (Lovett-Barron et al., 2014; Sadeh and Clopath, 2021). Alternatively, it is possible that these inhibitory motifs also exhibit plasticity during other forms of hippocampal learning. Future studies with combined molecular and physiological readouts in behaving animals are required to fully uncover both stable and plastic elements (Sadeh and Clopath, 2021; Vogels et al., 2011) in hippocampal inhibitory motifs.

## Acknowledgements

B.V. is supported by grants (NIH) T32GM007367 and (NIMH) F30MH125628. A.L. is supported by the National Institute of Mental Health (NIMH) 1R01MH124047, 1R01MH124867; National Institute of Neurological Disorders and Stroke (NINDS) 1U19NS104590 and 1U01NS115530, and the Kavli Foundation. The authors thank Zhenrui Liao and other members of the Losonczy lab for their invaluable comments on previous versions of the manuscript.

## Author contributions

All authors designed the project and experiments. B.V. and T.G. performed all experiments and analysis. All authors wrote the manuscript. T.G. and A.L. oversaw all aspects of the project.

## METHODS

### Animals

All experiments were conducted in accordance with NIH guidelines and with the approval of the Columbia University Institutional Animal Care and Use Committee. Experiments were performed with healthy, 3-5 month old, heterozygous adult male and female *VGAT-IRES-Cre* mice (Jackson Laboratory, Stock No: 016962) on a C57BL/6J background. Mice were kept in the vivarium on a reversed 12-hour light/dark cycle and housed 3-5 mice in each cage. Mice with implanted silicon probes were housed individually. Experiments were performed during the dark portion of the cycle.

### Viruses

Cre-dependent recombinant adeno-associated virus (rAAV) expressing GCaMP7f under the control of the Synapsin promoter (rAAV1-Syn-FLEX-jGCaMP7f-WPRE-Sv40, Addgene #104492, titer: 1 × 10^13^ vg/mL) was used to express GCaMP7f in interneurons.

### Virus injections and hippocampal window/headpost implant for CA3 imaging

For viral injections, 2 to 4-month-old *VGAT-Cre* mice were anesthetized with isoflurane and placed into a stereotaxic apparatus. Meloxicam and bupivacaine were administered subcutaneously to minimize discomfort. After the skin was cut in the midline to expose the skull, the skull was leveled, and a craniotomy was made over the right hippocampus using a drill. A sterile glass capillary loaded with rAAV1-Syn-FLEX-jGCaMP7f-WPRE-Sv40 was attached to a Nanoject syringe (Drummond Scientific) and slowly lowered into the right hippocampus. Dorsal CA3 was targeted with 2 X-Y coordinates, each one consisting of two different injection sites separated in Z: AP −1.35, ML −1.6, DV −2.1, −1.9 and AP −1.6, ML −1.9, DV −2.2, −2.0 relative to Bregma, with 50-64 nL of virus injected at each DV location. After injection, the pipette was left in place for 5-10 minutes and slowly retracted from the brain. The skin was closed with several sutures and the mice were allowed to recover for 4 days before the window/headpost implant.

The surgical procedure for CA3 window/headpost implant is very similar to the one implemented for CA1 imaging (Geiller et al., 2020). Briefly, the injected mice were anesthetized with isoflurane and placed into the stereotaxic apparatus. After subcutaneous administration of meloxicam and bupivacaine, the skull was exposed, leveled, and a 3 mm craniotomy was made over the right anterior hippocampus, centered between the two injection coordinates. The dura overlying the cortex was removed, and the cortex overlying the hippocampus was slowly removed with negative pressure while ice-cold cortex buffer was simultaneously applied. Care was taken to not damage the lateral ventricle. This process was performed until the white, anterior-posterior fibers overlying the hippocampus became visible and any bleeding subsided. A stainless steel, 3 mm wide X 2 mm long circular cannula fitted with a glass window was inserted into the craniotomy and pushed down to sufficiently flatten the natural curvature of the anterior hippocampus. The cannula was secured in place with Vetbond applied on the skull. Subsequently, dental cement was applied to the entire skull, and a headpost was affixed to the posterior skull with dental cement. The mice received a 1.0 mL subcutaneous injection of saline and recovered in their home cage while heat was applied. The mice were monitored for 3 days post-operatively until behavioral training began.

### AOD imaging

Prior to the random foraging experiments, mice first underwent a single imaging session consisting of a high-resolution structural scan. This step was necessary to obtain a reference Z-stack and derive the X-Y-Z positions of GCaMP-expressing neurons. The mice were head-fixed under a custom-modified AOD microscope (Femto3D-ATLAS, Femtonics Ltd) and anesthetized with ketamine/xylazine to reduce motion artifacts during the stack. To provide stable transmission parameters during chronic imaging in the entire 3D scanning volume, the AOD microscope was extended with a high speed and precision beam stabilization unit which was directly attached to the AOD scan head, sensitive to input beam misalignment. The beam stabilization unit consisted of two quadrant detectors (PDQ80A and TPA101, Thorlabs) and two broadband dielectric mirrors (Thorlabs) mounted on motorized mirror mounts (Femtonics). The beam alignment was performed by the LaserControl software (Femtonics). A water-immersion objective (16x Nikon CFI75) was placed above the glass window and lowered until the CA3 pyramidal cell layer was in focus. At this stage, the objective was fixed in position and focus was subsequently adjusted using AO crystals (Szalay et al., 2016). The laser (Coherent Ultra II) was tuned to λ=920 nm. The reference Z-stack was taken starting from the approximate location of CA2 (∼200-300 µm below the glass window) and extending as deep into CA3 as possible (∼500-600 µm below the glass window while still being able to visualize individual interneurons). 800×800 pixel images (X-Y resolution of 1.25 µm/pixel) were taken every 4. µm. Laser power and photomultiplier (PMT) detectors (GaAsP, H10770PA-40 Hamamatsu) were compensated appropriately in Z throughout the stack (power at 20-40 mW and detector gain at 80% at the top of the stack, power at 120-150 mW and detector gain at 90% at the bottom). After completion, the mice were returned to their home cage and allowed to recover for 24h until the start of functional imaging.

Prior to simultaneous imaging and LFP experiments, the mice were not anesthetized, and the reference Z stacks were taken on the same day as functional imaging was performed. Small Z stacks of ∼ 40um were taken while the mouse was immobile on the belt.

To determine X-Y-Z positions of GCaMP-expressing neurons, the Z-stack was scrolled through, and each visible interneuron was manually selected using the integrated software (MES, Femtonics Ltd) to generate a list of ∼100 X-Y-Z coordinates defined as the center of each cell (∼20 cells for simultaneous imaging/LFP experiments). These points constituted the centers of regions of interest (ROI) used on subsequent days for functional imaging. Each ROI was defined as a square of 40 to 50 µm^2^ (chessboard scan) (Szalay et al., 2016) with a resolution of 1 to 1.5 µm/px. The advantage of the chessboard scanning method is that only neurons and small areas around the pre-selected cells are recorded. Therefore, a high ratio of the total recording time (∼20-50%) is spent reading out information from the selected neurons. In contrast, volumetric imaging with the same 2P excitation provides an orders-of-magnitude worse ratio for measurement time utilization as the somata of interneurons occupy a relatively small ratio of the total scanning volume.

On each day of functional imaging, the same field of view was found using the reference Z-stack and X-Y-Z coordinates were loaded into the software to perform 3D imaging. Once all cells were in focus, 10 minute functional imaging sessions were conducted at a frame rate of 5-10 Hz for most experiments (frame rate was dependent on ROI size and resolution). For experiments involving contralateral LFP recordings, imaging was conducted at a higher rate (∼40 Hz), which restricted imaging to only 10-20 cells simultaneously. During functional imaging, the laser power and detector gain were compensated based on the reference Z-stack parameters.

### Silicon probe implantation, LFP recordings, and sharp-wave ripple identification

For experiments requiring simultaneous two-photon calcium imaging and LFP recordings, mice were implanted with a glass window over the hippocampus as above, and additionally a chronic, 4-channel silicon probe (Qtrode, Neuronexus) was inserted into the contralateral CA1 at a 45 degree angle. The probe was secured in place with dental acrylic and the mouse was allowed to recover for several days, as above. LFP signals were recorded with a multichannel recording system (Intan Technologies) synchronized with the AOD imaging system. The correct position of the silicon probe was confirmed by the presence of sharp-waves ripples in the data. LFP signals were recorded at 20kHz. To identify putative sharp-wave ripple events, the raw LFP signal was band-pass filtered from 150-300 Hz and thresholded at 2.5 standard deviations above the mean value within the passband. All putative sharp-wave ripple events were then manually inspected and false-positives were discarded to obtain the final set of sharp-wave ripple events used for analysis.

### Behavioral paradigm

After recovery from surgery, mice were handled for several days and habituated to head-fixation. Mice were subsequently water-restricted to 85-90% of their original weight and trained to run on a single-fabric, cue-free belt. Mice were trained to operantly lick and receive water rewards (water was delivered in response to tongue contact with a capacitive sensor) at random locations along the belt. As performance improved, the number of rewards delivered on each lap decreased. After several days of training on this cue-free belt, the mice were trained for ∼1 week on a 2m long, cue-rich belt for randomly delivered water rewards. For random foraging experiments, imaging was started after mice could run approximately 10 laps in 10 minutes (usually after 7-10 days of total training). For combined imaging and LFP experiments, data acquisition was started once GCaMP7f expression was optimal, hippocampal windows were clear, and the mice were habituated to head-fixation; these mice did not undergo additional behavioral training. The sensory stimulation experiments (random cue task) were performed as described previously (Terada et al., 2021) on a burlap belt. Briefly, three sensory cues (odor, light, and a non-operant water reward) were presented randomly at 15 trials per cue independently of the mouse’s position on the treadmill and each cue presentation was separated by a random inter-stimulus interval of 10-15s.

### Perfusion and tissue processing

After the completion of imaging experiments, mice were transcardially perfused with 40 mL of ice-cold Phosphate-Buffered Saline (PBS, Thermo Fisher), followed by 40 mL of ice-cold 4% paraformaldehyde (PFA, Electron Microscopy Sciences). Brains were stored overnight in 4% PFA at 4°C. The next day, the 4% PFA was removed and the brains were rinsed 3×5 min in PBS. 75 µm horizontal sections of the imaged hippocampus were cut on a vibrating microtome (Leica VT1200S) and washed 3×15 minutes in PBS. Subsequently, sections were permeabilized for 2×20 minutes in PBS with 0.3% Triton X- 100 (Sigma-Aldrich). Blocking was then performed with 10% Normal Donkey Serum (Jackson ImmunoResearch, Catalog #017-000-121) in PBST (PBS with 0.3% Triton X- 100) for 45 minutes. The sections were then incubated in a PBS solution containing 3 primary antibodies (see below for antibody information and dilutions) for one hour at room temperature, followed by 2 days at 4°C. After 2 days, the primary antibody solution was removed from the slices and the slices were washed 3×15 minutes in PBS to remove unbound primary antibodies. The slices were subsequently incubated in a PBS solution containing a mixture of appropriate secondary antibodies conjugated to fluorescent labels (see below for antibody information and dilutions) for 2 hours at room temperature. The sections were then washed 5×15 minutes in PBS at room temperature. Finally, sections were mounted on glass slides in Fluoromount-G aqueous mounting medium (ThermoFisher Scientific) and coverslipped. The slides were allowed to dry at 4°C for at least one hour before confocal imaging (see below). After confocal imaging, the slides were submerged in PBS to remove the coverslip, and the sections were removed from the slides with gentle rocking. After washing 3×15 min in PBS and blocking with 10% Normal Donkey Serum in PBST for 45 minutes, the sections were incubated in an additional 2-3 primary antibodies. The sections were subsequently washed, incubated in secondary antibodies, washed again, and mounted and imaged, as in the first round of staining.

### Immunohistochemistry (See Resources table for antibody catalog numbers)

#### Random foraging mice (Figure 2 data)

First round primary antibodies: rabbit anti-proCCK (1:500), rat anti-somatostatin (1:500), and goat anti-calbindin (1:500)

First round secondary antibodies: donkey anti-rabbit DyLight 405 (1:300), donkey anti-rat Rhodamine Red (1:300), and donkey anti-goat Alexa 647 (1:300)

Second round primary antibodies: chicken anti-PV (1:5,000) and rabbit anti-SATB1 (1:1,000)

Second round secondary antibodies: donkey anti-chicken DyLight 405 (1:300) and donkey anti-rabbit Rhodamine Red (1:300)

#### SWR mice(Figures 3-5 data)

First round primary antibodies: chicken anti-PV (1:5,000), rat anti-somatostatin (1:500), and rabbit anti-SATB1 (1:1,000)

First round secondary antibodies: donkey anti-chicken DyLight 405 (1:300), donkey anti-rat Rhodamine Red (1:300), and donkey anti-rabbit Alexa 647 (1:300)

Second round primary antibodies: rabbit anti-proCCK (1:500) and goat anti-calbindin (1:500)

Second round secondary antibodies: donkey anti-rabbit rhodamine red (1:300) and donkey anti-goat Alexa 647 (1:300)

### Confocal imaging

Confocal imaging: A Nikon A1 confocal microscope was used to acquire multi-channel fluorescence images of the immunolabeled tissue sections. 405 nm, 488 nm, 561 nm, and 640 nm laser lines were used for excitation. Each channel was acquired sequentially with a 10x Plan Apo NA 0.45 objective (Nikon) at 1.2-1.3x Zoom. 2048x 2048 pixel images were acquired every ∼3 microns through the entire depth of the tissue sections, with the pinhole size set to ∼1 Airy unit. Fluorescence was collected with 2 GaAsP PMTs (488 nm and 561 nm channels) and 2 multi-alkali PMTs (405 nm and 640 nm channels). The resulting 4-channel Z-stacks were viewed in ImageJ (NIH).

### Registration of confocal images to *in vivo* Z-stacks and identification of immunopositivity/negativity

The following steps were performed by an experimenter without the use of any automated methods: First, the confocal stacks were rotated and translated until the cells in the green channel (GCaMP-labeled) matched the cells seen in the in vivo Z stack. Second, each imaged cell was found in the confocal stacks, and it was evaluated for immunopositivity or immunonegativity for the tested molecule. For a cell to be considered positive, the fluorescence intensity inside the cell had to be significantly greater than the background intensity level. A cell was considered positive for a given marker only if clear examples of immunonegative cells could be found on the same tissue section. Similarly, a cell was considered negative for a given marker only if clear examples of immunopositive cells could be found on the same tissue section. In the case of ambiguous immunolabeling, cells were discarded and not grouped into a subtype for further analysis. Overall, all efforts were made to use the most stringent criteria for cell classification prior to analysis.

### Subtype assignment

Subtypes were assigned based on the immunoreactivity of cells to the five tested markers and the association between these markers and defined interneuron subtypes, based on the previous literature. All imaged cells not within the region innervated by CB-positive mossy fibers were assumed to be within CA1 and were excluded from all analyses.

**PVBC:** PVBCs were positive for PV and SATB1, and negative for the other three tested markers (SOM, CCK, and CB).

**AAC:** AACs were positive for PV but negative for SATB1, and negative for the other three tested markers (SOM, CCK, and CB).

**SOM:** All cells positive for Somatostatin were categorized as SOM cells. While all cells within this category were negative for CCK, some were also positive for PV, SATB1, or CB. Notably, Somatostatin/Calbindin co-expressing cells were included in this category because they represent long-range projecting interneurons located almost exclusively in *stratum oriens* of CA1-CA3 (Jinno, 2009; Jinno et al., 2007). Thus, the SOM subtype here represents putative dendrite-targeting and some long-range projecting interneurons.

**CCK:** CCKs were necessarily positive for CCK, and all of these cells were always negative for PV and Somatostatin. Cells within this category could express SATB1, although the vast majority of them were SATB1-negative. Although some cells within the dataset co-expressed CCK and Calbindin, the vast majority of CCK- and/or CB- expressing cells expressed only one of the two markers. Thus, CCK and Calbindin double-positive neurons were excluded from further analysis, and this category represents those cells positive for CCK only.

**CB:** CBs were necessarily positive for CB, and all of these cells were always negative for PV. Although some cells co-expressed Somatostatin and Calbindin, these cells were included in the SOM subtype (see above). Similarly, while some cells co-expressed CCK and CB, only the CB-positive and CCK-negative cells are included in this subtype (see above). CB cells could be either positive or negative for SATB1. Thus, this category represents putative dendrite-targeting, Calbindin-expressing interneurons (Gulyás and Freund, 1996).

### Calcium imaging data preprocessing

The raw movies containing each cell were motion corrected independently using a whole-frame cross-correlation algorithm, as implemented in the SIMA software (Kaifosh et al., 2014). The time-average of each imaged cell was manually inspected and a ROI was hand-drawn over each cell. Fluorescence was extracted from each ROI using the FISSA software (Keemink et al., 2018) package to correct for neuropil contamination, using 6 patches of 50% the size of the original ROI. For each resulting raw fluorescence trace, a baseline F was calculated by taking the 1^st^ percentile in a rolling window of 30s and a Δ*F/F* trace was calculated. The Δ*F/F* trace for each cell was smoothed using an exponential filter and all further analysis was performed on the resulting *ΔF/F* traces. All further analyses were implemented in Python 2.7 and are detailed below.

### Locomotion and immobility modulation

To calculate the correlation between each cell’s activity and the animal’s velocity, the Pearson’s correlation coefficient was calculated between each cell’s Δ*F/F* trace and the smoothed velocity trace.

### Run-start and run-stop responses

Run-start and run-stop events were identified in the imaging data as frames during which the animal’s velocity increased above 0.2 cm/s (run-start event) or decreased below 0.2 cm/s (run-stop event). In addition, each run-start/run-stop event had to be separated from the previous run-start/run-stop event by at least several seconds to be considered as a separate event. For each event, the mean of the pre-event Δ*F/F* was subtracted from the mean of the post-event Δ*F/F* in a −3s to +3s window to calculate a response magnitude. For each cell, the run-start and run-stop response magnitudes were averaged over all run-start and run-stop events in the given imaging experiment. If a cell was imaged across more than one imaging experiment, the average run-start and run-stop responses from each experiment were averaged over all experiments.

### Spatial tuning curves

To calculate a spatial tuning curve for each imaged cell in a given experiment, the 2m treadmill was divided into 100 2cm-long bins. For each bin, we calculated the average Δ*F/F* from frames where the animal was in locomotion (velocity > 5cm/s). To determine whether a cell was spatially tuned during an imaging session, we generated 1,000 shuffled tuning curves by circularly rotating position in relation to Δ*F/F* traces (restricted to frames during locomotion). A cell was detected spatially selective if it had 10 consecutive bins (20cm) exceeding the 95^th^ percentile of the shuffle distribution (or lower than the 5^th^ percentile for negative fields). Place field centroids were calculated as the center of mass of the cell’s tuning curve.

### Generalized linear model

To more explicitly dissociate the effects of the various behavioral variables on each cell’s activity during navigation, we developed a multivariate linear regression model to predict each cell’s fluorescence activity (ΔF/F) from the following behavioral variables: 1) the animal’s velocity, 2) position, 3) reward, and 4) licking. The position variable was itself divided into 10 predictors, corresponding to 10×20cm segments of the treadmill. The model utilized Ridge regression to minimize the effects of potential relationships between the independent variables. The fit values (R^2^) for full and reduced (lacking a given predictor) models are cross-validated with a leave-one-out procedure where 1 lap is left out in the time domain.

### Peri-SWR time histogram

To construct the average peri-SWR time histogram for a given cell, the cell’s ΔF/F trace was Z-scored in a −3s to +3s window around each SWR event, and the resulting peri-SWR traces were averaged together across all SWR events to obtain one trace. All SWR events were considered; none were excluded.

### PCA

Decomposition of peri-SWR time histograms, hereafter referred to as PSTHs, was performed using Principal Component Analysis in the scikit-learn library. Each interneuron’s average PSTH was projected onto the two first principal components and the median loadings for each subtype were calculated in PC space.

### SWR activity index

For each cell, a baseline activity was calculated as the mean fluorescence activity 3s to 2s prior to a given SWR (−3 to −2s from onset) from the cell’s average PSTH. Then, both a negative and positive modulation value was computed by respectively subtracting the baseline from the absolute minimum and maximum activity value in the window −1s to +1s from onset. The largest value of the two was kept. If the modulation value was derived from the maximum, the cell was considered activated; otherwise, the cell was considered inhibited.

### SWR power

To obtain the amplitude of each SWR event, the broadband LFP trace between the start and end of each detected SWR event was band-pass filtered between 150-300 Hz, and the maximum of the absolute value of the filtered trace was taken as the amplitude. To compare a given cell’s activity across SWR events of varying power, all SWRs which occurred during the time that cell was imaged were Z-scored. Thus, while SWR power is highly dependent on the electrode position within CA1, all comparisons were made only between SWR events that occurred on the same day in a given mouse. Additionally, because we consider only Z-scored SWR amplitudes and not absolute figures, we use the terms ‘SWR amplitude’ and ‘SWR power’ interchangeably.

To calculate PSTHs for each subtype around low- and high-power SWRs, we first calculated PSTHs around low (0-20^th^ percentile) and high (80^th^-100 percentile) power SWRs for each cell. The PSTHs for all cells within a subtype were then averaged. To calculate a difference value between high- and low-power SWRs for each cell, the cell’s low-power PSTH was subtracted from its high-power PSTH, and the average value of this difference curve in a 500 ms window around the SWR (−500 ms to + 500 ms) was taken as the cell’s difference value. The location of the maximum difference between the high- and low-power PSTHs within this window was taken as the maximum difference.

### Reflective/predictive activity

To correlate the pre-SWR activity of each cell with SWR power, the cell’s ΔF/F trace was first Z-scored in a −3s to +3s window around each SWR event, as above. Then, for each SWR, the cell’s average ΔF/F value in the 500ms before the event and the event’s power were considered. These two sets of values were fit with a linear regression model, and the R value of the regression was recorded. To minimize spurious correlations, only cells recorded during at least 100 SWR events were considered; the vast majority of cells were recorded for 500-1,500 SWR events. The same procedure was used to correlate the post-SWR activity of each cell with SWR power, but now the cell’s average ΔF/F value in the 500 ms following each SWR event was considered.

### Transient detection of CA3 pyramidal cells

Transient intervals were detected from the CA3PC data as intervals where the Z-scored activity for a given cell exceeded 2 and stayed above 0.5 for at least 0.5 seconds. The first frame of this interval was taken as the transient onset and was used for analysis.

### Cue responses

The cue response for each interneuron was calculated as the difference between the mean activity from 0 to 1s after cue presentation and the baseline (−1 to 0s before), regardless of cue identity. Cue-specific responses for each subtype are reported in the Supplementary Information.

### SWR modulation

To calculate the SWR modulation for a given cell, the average peri-SWR time histogram was first computed, as described above. We defined the SWR modulation as the maximum of this average trace in the 500-ms interval following the SWR event minus the average of the pre-SWR baseline (defined as the −3 to −1 second interval preceding the SWR). To compare the SWR modulation values for a given cell between the PRE and POST sessions, the cell must have been imaged during at least 100 SWRs in both the PRE and POST sessions.

### SWR recruitment

A given cell’s SWR recruitment was calculated as the fraction of SWR events during which its response exceeded the 95^th^ percentile of a shuffle distribution. The cell’s response to an individual SWR was taken as the maximum of its activity within the 500-ms interval following each SWR, and the shuffle distribution was created by repeating this calculation for 1,000 randomly selected frames that occurred during immobility. To compare the SWR recruitment values for a given cell between the PRE and POST sessions, the cell must have been imaged during at least 100 SWRs in both the PRE and POST sessions.

### Statistical analysis

Statistical details of comparisons are specified in either the main text or figure legends. No statistical methods were used to predetermine sample sizes, but our sample sizes are similar to those reported in previous studies. Box plots represent median and interquartile range (IQR) while whiskers extend to cover the distribution without outliers (defined as points above 1.5 IQR below or above the box edges). Bar plots represent mean and s.e.m. Between-subtype comparisons were tested using a one-way ANOVA followed by a Tukey’s range test with correction for multiple testing if appropriate. For comparisons between two populations, a paired sample or unpaired t-test was applied if the data points followed a normal distribution. To analyze data that was not normally distributed, the Mann-Whitney U test was used. *, p < 0.05, **, p < 0.01, ***, p < 0.001. Data analysis and figures were done using custom made software in Python 2.7.15^TM^.

## SUPPLEMENTARY FIGURE LEGENDS

**Supplemental Figure 1.**
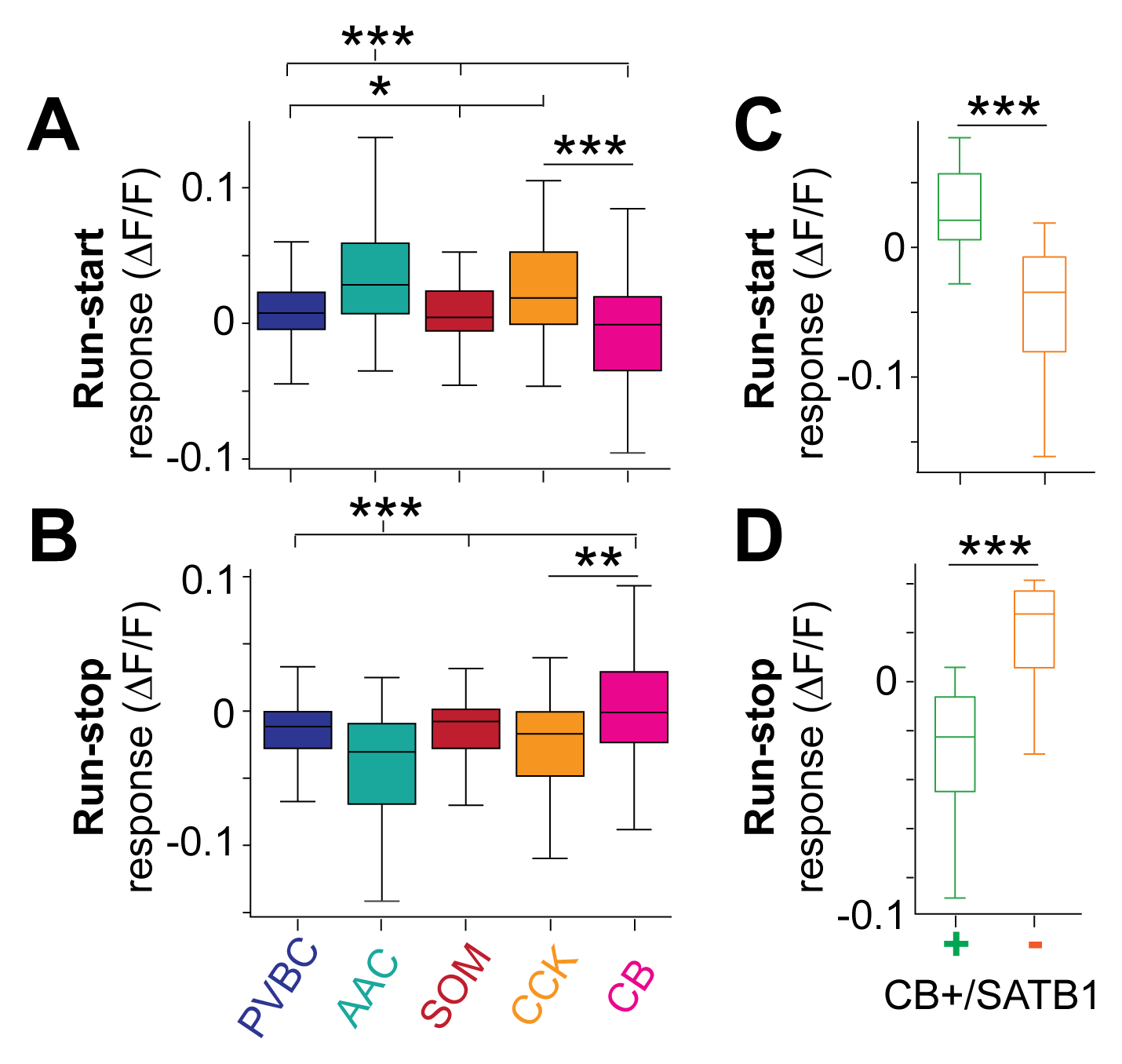
Quantification of locomotory response onsets. A. Quantification of run-start responses for all main subtypes (see Methods). AACs and CCKs had larger responses than many other subtypes (one-way ANOVA with post-hoc Tukey’s range test). B. Same as A, but for run-stop responses (see Methods). C. Same as A, but now splitting CB interneurons according to their immunoreactivity for SATB1. CB+/SATB1+ neurons had significantly larger run-start responses than CB+/SATB1- neurons (Mann-Whitney U Test). D. Same as C, but for run-stop responses.

**Supplemental Figure 2.**
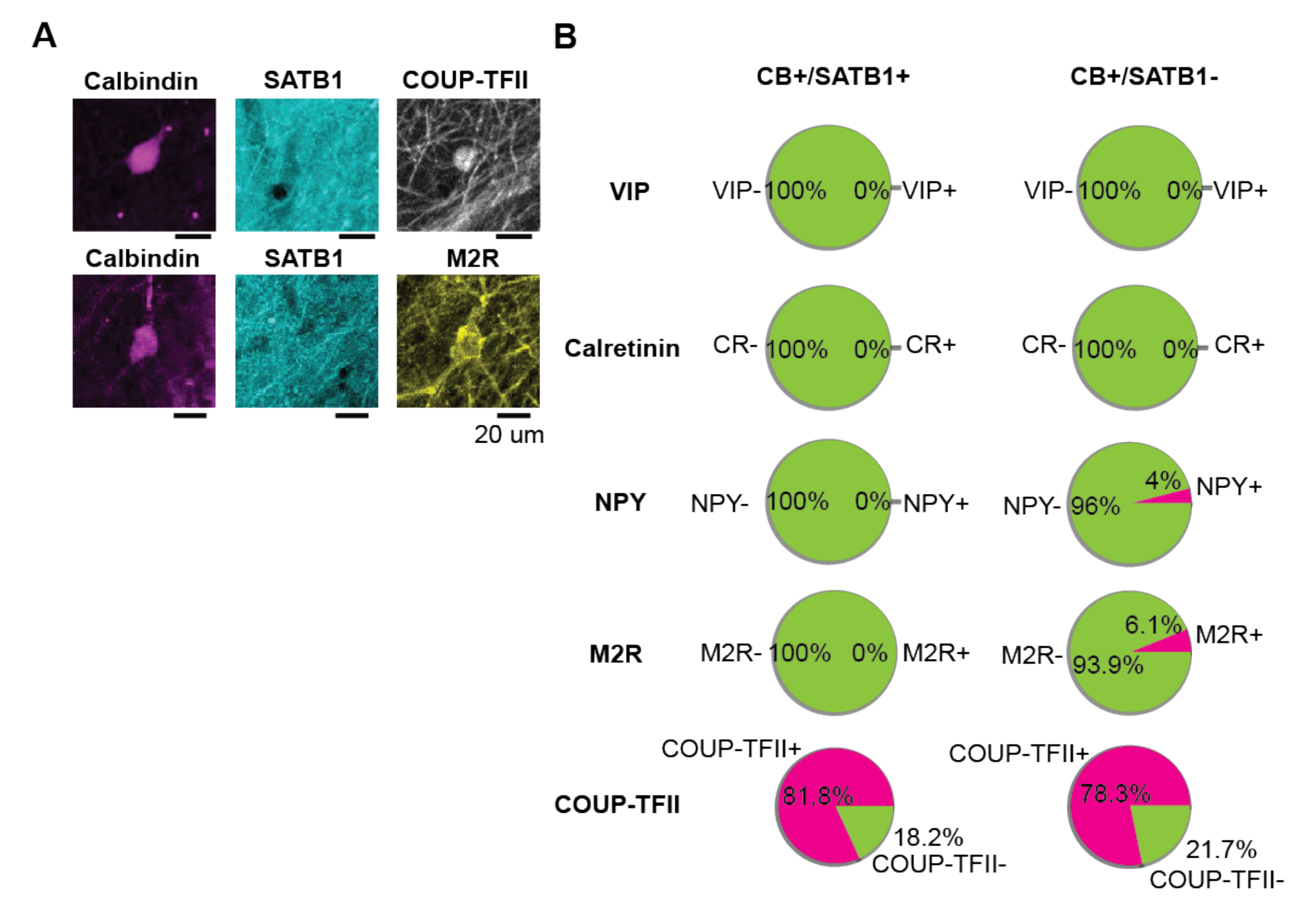
Molecular profiling of calbindin-positive SATB1-negative immobility-active interneurons. A. Confocal micrograph of CB-expressing interneurons, negative for SATB1 but positive for COUP-TFII (top) and M2R (bottom). Scale bars represent 20 µm. B. Quantification of the overlap of CB-expressing interneurons split by immunoreactivity to SATB1 with other markers.

**Supplementary Figure 3.**
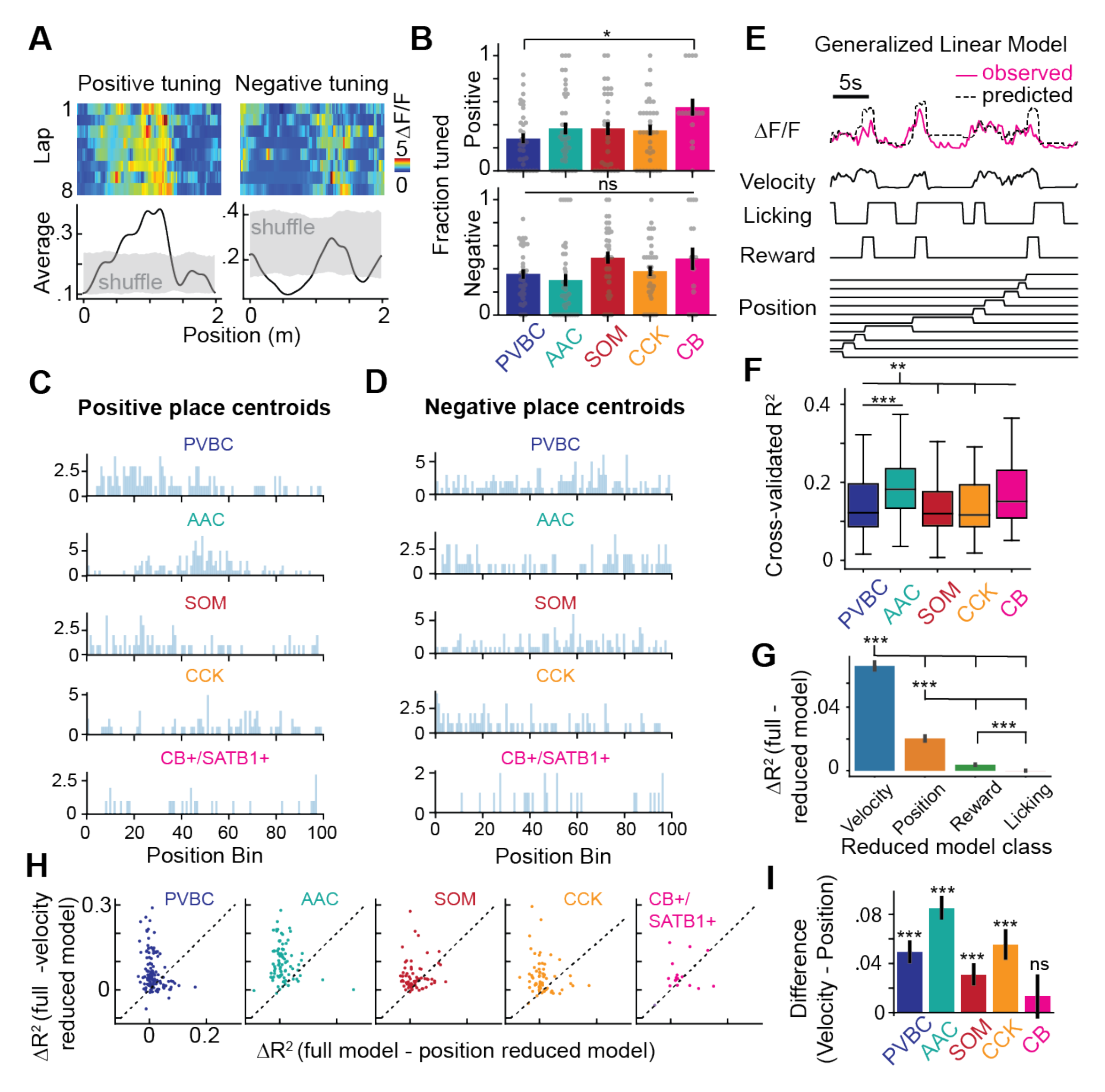
Interneuron spatial selectivity during navigation. A. Representative examples of spatially-selective interneurons with positive (left) and negative tuning (right), defined based on a shuffled distribution (see Methods). B. Fractions of positively (top) and negatively (bottom) tuned cells by subtype. Each dot represents the fraction of tuned cells for a given subtype during a recording session. CBs had significantly greater fractions of tuned cells than did PVBCs (one-way ANOVA with *post-hoc* Tukey’s range test). Note the CB label here represents only CB+/SATB1+ neurons, as CB+/SATB1- cells were shown to be silent during locomotion. C. Distribution on the belt of all positive place field centroids. D. Same C, but for negative place field centroids. E. Schematic of the GLM used to evaluate the contribution of multiple predictors to each interneuron’s fluorescence activity. F. Distribution of goodness of fit (R^2^) between the predicted and actual observed activity for each subtype. AACs had significantly greater R^2^ values than most other subtypes (one-way ANOVA with *post-hoc* Tukey’s range test). G. Difference in R^2^ between the full model (with all predictors from E) and various reduced models, each one lacking one predictor. Each category of reduced model was fit over all cells, regardless of subtype. The velocity predictor contributed most to activity, followed by position, then reward, and finally licking (paired t-tests with *post-hoc* Bonferroni correction). H. Contribution of position (X-axis) versus velocity (Y-axis) to the activity of each cell, measured as the difference in R^2^ in reduced models lacking these predictors. Each dot represents an interneuron. The activity of some cells can be better explained by either one of the two predictors. I. Difference between the contributions of velocity and position to the activity of each cell shown in H) for each subtype. For most subtypes, the velocity predictor could explain activity better than the position predictor (one-sample t-tests against 0).

**Supplementary Figure 4.**
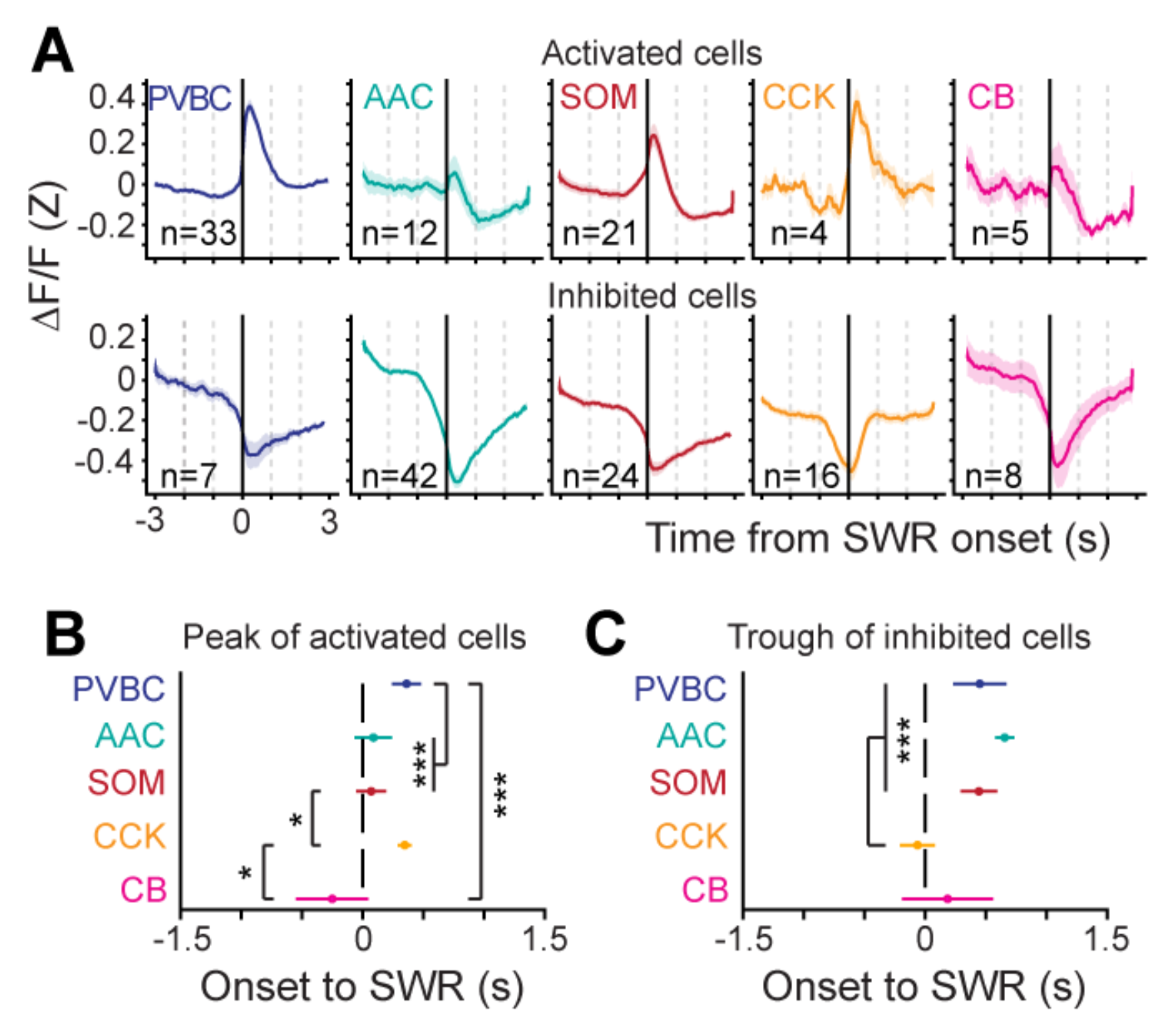
Subtype-specific inhibited and activated response dynamics to SWRs. A. Average peri-SWR time histogram for activated (top) and inhibited (bottom) interneurons, defined by the SWR modulation index (see Methods). The number of activated and inhibited cells for each subtype is reported in *Figure 3*. B. Timing of the peak location computed for individual activated neurons for each subtype (mean ± sem). C. Same as B, for the trough location of inhibited neurons (mean ± sem).

**Supplementary Figure 5.**
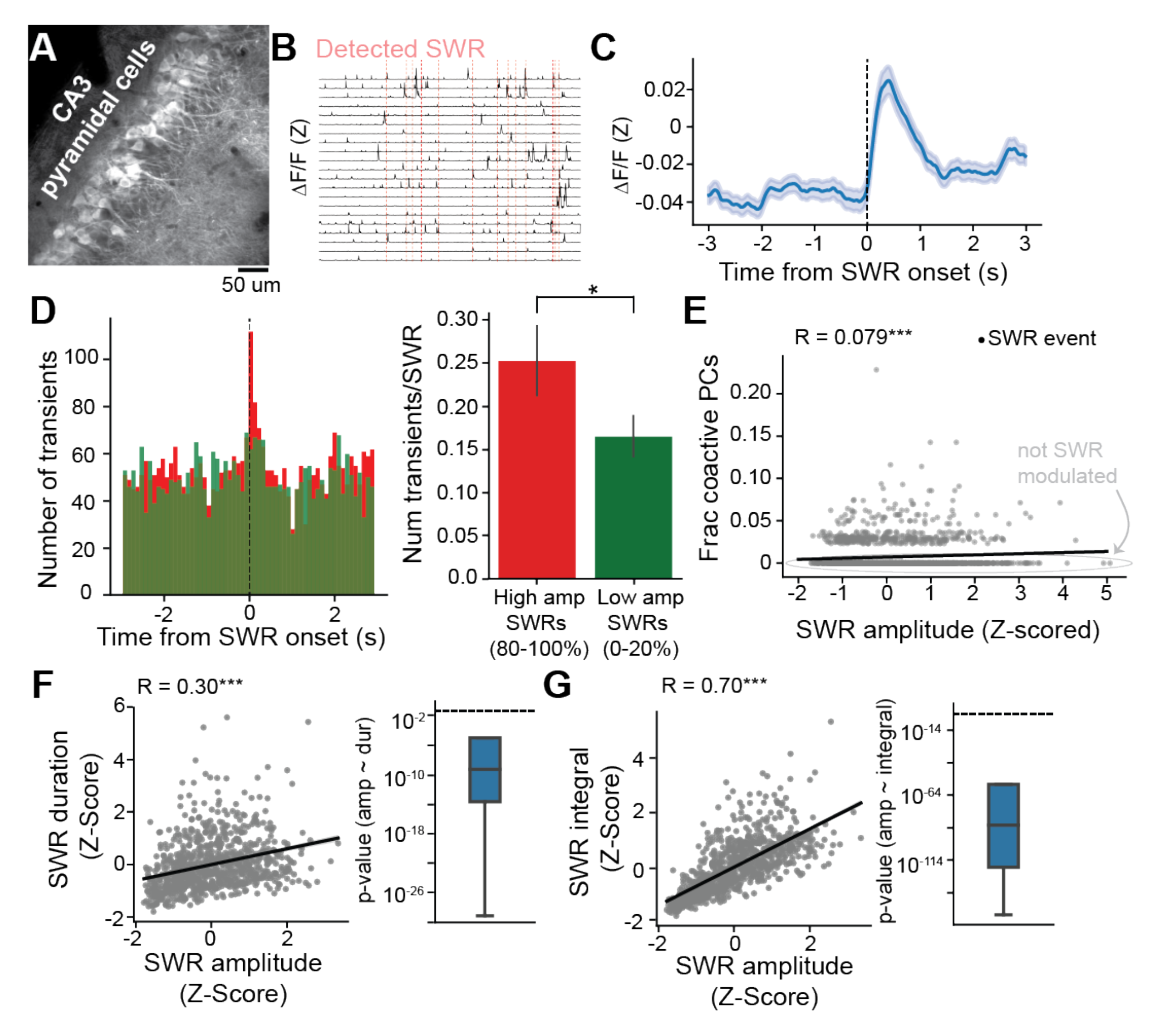
CA3PC data and Correlations between SWR Properties. A. Representative *in vivo* two-photon time-average image of a CA3PC FOV. B. Example CA3PC ΔF/F traces with detected SWRs depicted as vertical red lines. C. Peri-SWR fluorescence activity for all CA3PCs, averaged over all SWR events. D. *Left*: Distribution of peri-SWR CA3PC calcium transients for SWRs with low power (green, taken as SWRs with power falling between 0-20^th^ percentile of all ripples for a given mouse) and high power (red, for SWRs between 80-100^th^ percentile). *Right*: Quantification of the population transient rate for high- and low-power SWRs. CA3PCs emitted significantly more transients during high-power SWRs than during low-power SWRs (Wilcoxon signed-rank test). E. Correlation between SWR power and the number of co-active pyramidal cells around the SWR. High-power SWRs were associated with greater fractions of co-active CA3PCs around the SWR event. F. Correlation between amplitude and duration for individual SWRs. *Left:* Example scatter plot and linear regression line depicting the relationship between amplitude and duration for all SWRs recorded during the imaging of one example interneuron. *Right*: Distribution of p-values for the regression between amplitude and duration, calculated over all imaging sessions. The horizontal dashed line corresponds to a p-values of 0.05. A strong relationship between SWR amplitude and duration was present in all imaging sessions. G. Same as F, but now performed for the relationship between SWR amplitude and SWR integral. Again, these two LFP measures were strongly correlated.

**Supplementary Figure 6.**
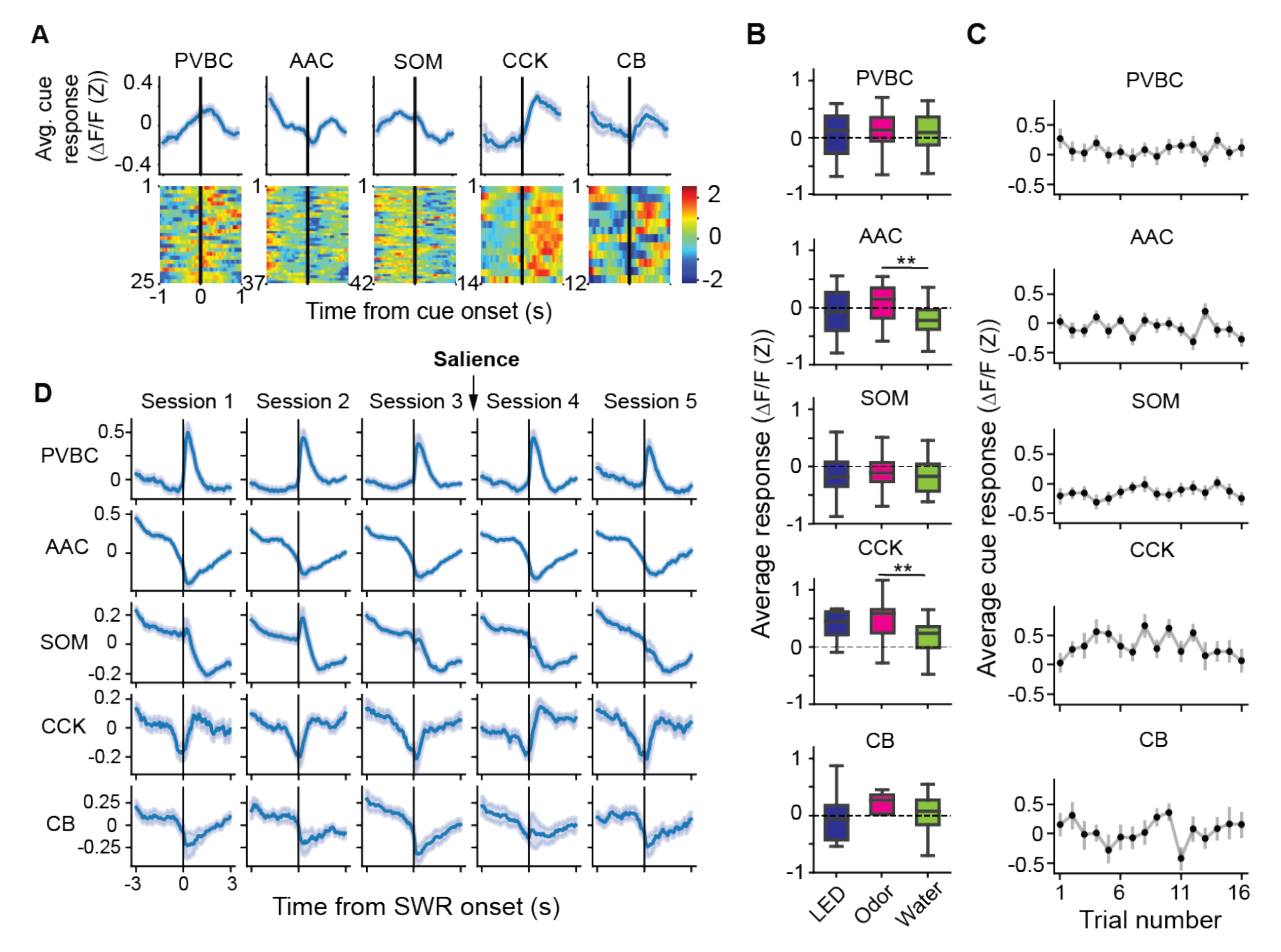
Session-to-session and trial-to-trial cue responses and SWR-associated dynamics in the sensory stimulation task. A. Average cue response traces for each subtype (top) and the response for each individual neuron (bottom; each row represents one cell). B. Average cue response for each subtype, broken down by the response to each sensory modality (visual stimulation, odor delivery, and water delivery). While most subtypes responded similarly to the three sensory cues, AACs and CCKs exhibited greater responses to odor than to water stimuli (paired t-tests with *post-hoc* Bonferroni correction; only significant comparisons are shown). C. Average cue response across trials for each subtype. Each trial represents the average of three sensory stimulations, one of each subtype (one visual, one odor, and one water stimulation). D. Peri-SWR ΔF/F traces for each session and subtype. Sensory stimulation occurred between Session 3 and Session 4. In Main Figure 5, Sessions 1-3 are considered PRE and Session 4 is considered POST.

